# Covering All Bases: A Computational Method to Design Broad-spectrum T-cell-inducing Vaccines Applied to *Betacoronaviruses*

**DOI:** 10.1101/2025.04.14.648815

**Authors:** Phil Palmer, Sofiya Fedosyuk, Srivatsan Parthasarathy, Jonathan Holbrook, Charlotte George, Laura O’Reilly, Lara Wiegand, George William Carnell, Jonathan Luke Heeney, Sneha Vishwanath

## Abstract

Antigenically diverse pathogens, such as coronaviruses, pose substantial threats to global health. This highlights the need for effective broad-spectrum vaccines that elicit robust immune responses in a large proportion of the human population against a wide array of pathogen variants. Here, we introduce Spectravax, an AI-enabled computational method to design broad-spectrum vaccines that account for genetic diversity in both the host and pathogen populations. Using Spectravax, we designed a nucleocapsid (N) antigen to elicit cross-reactive immune responses to viruses from the *Sarbecovirus* and *Merbecovirus* subgenera of *Betacoronaviruses*. *In silico* analyses demonstrated superior predicted host and pathogen coverage for Spectravax compared to wild-type sequences and existing computational designs. Experimental validation in mice supported these predictions: Spectravax N elicited robust immune responses to SARS-CoV, SARS-CoV-2, and MERS-CoV—the three coronaviruses responsible for major outbreaks in humans since 2002—while wild-type and existing computational designs elicited limited responses. Furthermore, we were able to identify the MERS-CoV N epitopes responsible for Spectravax’s cross-reactivity. Thus, we advance the rational design of broad-spectrum vaccines for pandemic preparedness.

## Main

Antigenically diverse and rapidly evolving pathogens, such as coronaviruses, pose substantial challenges to global health. Vaccines are among the most cost-effective medical interventions and are estimated to have saved 20 million lives from the Severe Acute Respiratory Syndrome Coronavirus 2 (SARS-CoV-2) pandemic in 2021 alone^1^. However, most of the available vaccines are designed to target a single strain or closely related strains. This limitation reduced their effectiveness against emerging variants, such as Omicron and its sublineages, which evaded immune responses generated by prior infection, vaccination or both^2^. Additionally, other betacoronaviruses (*β*-CoVs), including viruses from the *Sarbecovirus* and *Merbecovirus* subgenera, pose risks of zoonotic spillover^3^. This emphasises the need for broad-spectrum vaccines that provide protection across multiple strains and variants.

According to recent estimates, a broadly protective coronavirus vaccine could have prevented 40–65% of COVID-19 deaths during the pandemic’s first year^4^ and could be worth at least $1.5 trillion in social value to the US alone^5^. Calls for a pan-*β*-CoV vaccine aim to prevent diseases caused by SARS-CoV, SARS-CoV-2, Middle East Respiratory Syndrome Coronavirus (MERS-CoV), and future emergent coronaviruses^6^. In response, substantial investments have been made toward developing pan-coronavirus vaccines^7^. However, most pan-*β*-CoV vaccines in development use multivalent approaches, such as nanoparticles displaying multiple antigens from different viruses^8;9^, which may face manufacturing and regulatory challenges due to their complexity^10^.

Computational methods offer a generalisable solution for designing single-antigen broad-spectrum vaccines as they can easily be applied to other pathogens^11^. These computationally designed antigens can serve as alternatives to, or components of, multivalent vaccines, enhancing their breadth^12^. Approaches like Epigraph^13^ generate mosaic antigens that maximise coverage of pathogen sequence diversity. Deep learning-based immunoinformatics tools such as NetMHCpan^14^ predict peptide-MHC binding affinities, enabling the selection of epitopes that are more likely to elicit T-cell responses. OptiVax^15^ combines these predictions with human leukocyte antigen (HLA) haplotype frequencies to optimise peptide-based vaccine designs for host population coverage. However, few methods integrate pathogen diversity and host immunogenicity to design conserved and immunogenic vaccines^15;16;17^, and none integrate both host and pathogen coverage to maximise the number of individuals protected against the greatest number of pathogen variants.

Here, we focus on predicting T-cell epitopes, which is currently more tractable than predicting B-cell epitopes due to their simpler, short linear peptide sequences, as opposed to the complex three-dimensional structures critical for B-cell epitope recognition^14;18;19^. Focusing on T-cell-mediated immunity offers additional benefits. T-cell-mediated immunity is crucial for controlling viral infections and provides long-lasting protection^20^. T-cells can recognise conserved internal viral proteins less susceptible to mutation, offering broader protection against diverse strains^19^. In coronaviruses, robust T-cell responses have been associated with effective clearance of infections^21;22^, milder disease^23^, reduced transmission^24^, long-lasting immunity^25^, and decreased susceptibility to immune escape^26^. The nucleocapsid (N) protein is highly conserved, immunogenic and N-specific T-cells are correlated with protection against SARS-CoV-2 infection^27^, making it an ideal candidate for broad-spectrum vaccine design^28^.

In this work, we present Spectravax, a computational framework and method to design single-antigen broad-spectrum vaccines optimised to account for genetic diversity in both pathogen antigens and host Major Histocompatibility Complexes (MHCs). Spectravax leverages peptide-MHC binding predictions and host HLA allele frequencies to select antigen sequences predicted to be widely recognised by T-cells across diverse populations, while using k-mer frequencies to ensure conservation across pathogen strains. We applied this method to design an N antigen (Spectravax N) intended to elicit cross-reactive immune responses to viruses from the *Sarbecovirus* and *Merbecovirus* subgenera. Our analyses show that Spectravax N is predicted to display a wide range of T-cell epitopes across a diverse set of HLA alleles, providing robust host population coverage across three different ancestries and 95 countries. Furthermore, Spectravax N has high sequence similarity to N proteins from *Sarbecoviruses* and *Merbecoviruses* and is the only vaccine candidate in our study predicted to induce T-cell responses to all target pathogen sequences, when compared to wild-type sequences and a vaccine design generated using Epigraph^13^. We chose to compare our design with Epigraph, as our method uses a similar graph-based algorithm for vaccine design. Experimental validation in a mouse model demonstrated that Spectravax N elicited immune responses against all three tested coronaviruses, while the wild-type and Epigraph antigens induced narrower responses. Finally, using smaller peptide pools from MERS-CoV N for ELISpot and flow cytometry assays, we were able to identify the specific epitopes responsible for Spectravax N’s MERS-CoV reactivity. Taken together, these findings highlight Spectravax as a promising approach for the rational design of broad-spectrum vaccines.

## Results

### Target antigen selection

We selected the Nucleocapsid (N) protein as the target antigen based on its high T-cell immunogenicity and sequence conservation across the *Sarbecovirus* and *Merbecovirus* subgenera (Fig. 1).

**Fig. 1:**
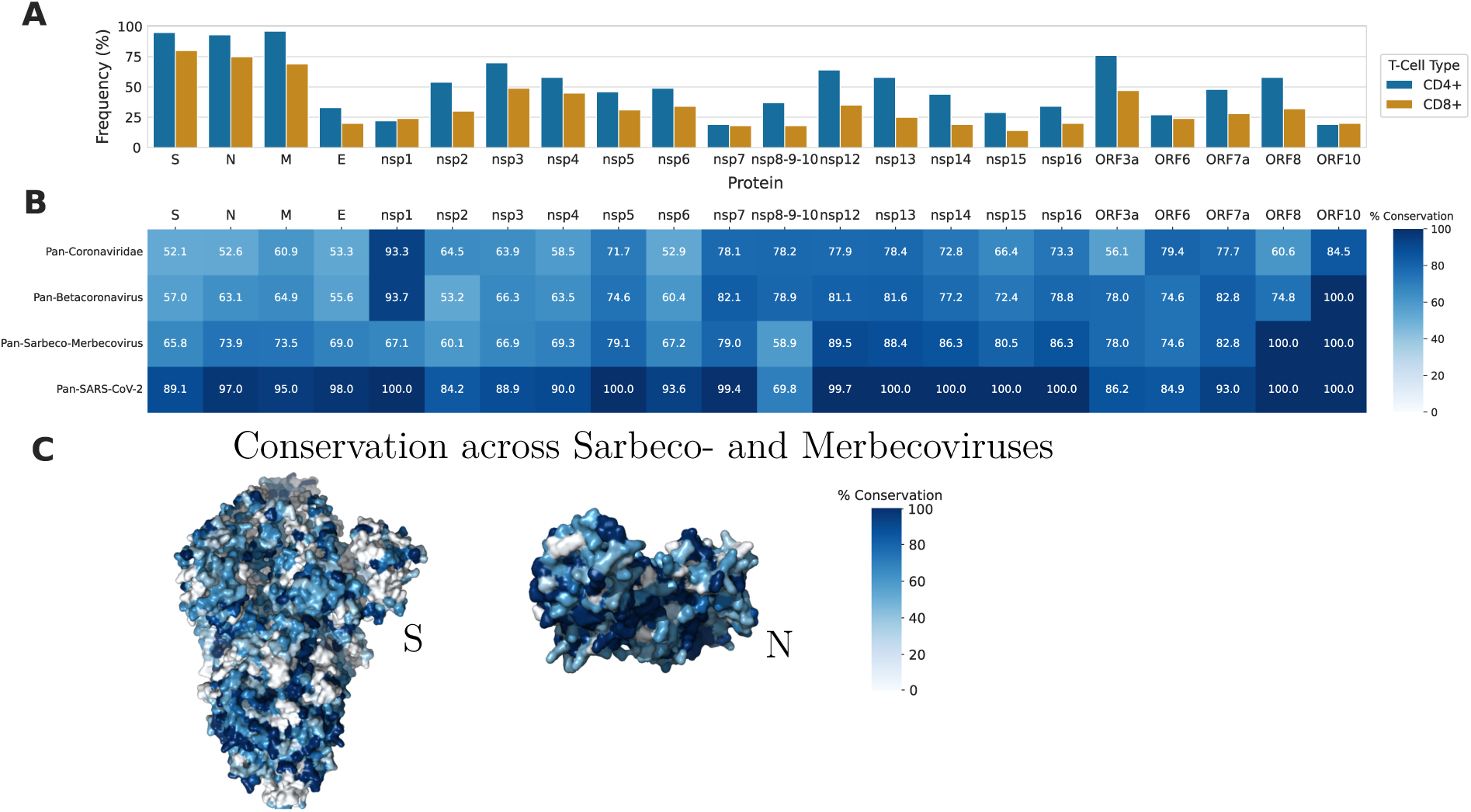
Conservation and T-cell immunogenicity of *Coronaviridae* proteins. **A)** Reported CD4^+^ and CD8^+^ immunogenicity of SARS-CoV-2 proteins in convalescent COVID-19 donors (*n* = 99)^29^. **B)** Protein sequence conservation of coronavirus proteins across the *Coronaviridae* family, *Betacoronavirus* genus, *Sarbecovirus* and *Merbecovirus* subgenera, and SARS-CoV-2 strain. All *Coronaviridae* proteins with full RefSeq genome completeness were downloaded from the NCBI Virus database^30^ (accessed on 25th June 2024) and aligned using MAFFT^31^. Sequence conservation was calculated for each protein as the mean percentage identity to the most common amino acid at each position, excluding gaps. **C)** Surface representation of the spike (S) ectodomain trimer (PDB ID: 6VSB^32^) and nucleocapsid (N) single subunit (PDB ID: 8FG2^33^) of SARS-CoV-2. The amino acids are color coded according to the sequence conservation across the *Sarbecovirus* and *Merbecovirus* subgenera. The colour bar represents the percentage of conservation. The image was generated and rendered using Pymol Open-Source v2.5.0^34^.

The N protein has the second-highest total T-cell immunogenicity among SARS-CoV-2 proteins, only slightly below the Spike (S) protein (Fig. 1A). Specifically, 93% and 75% of individuals responded to the N protein for CD4^+^ and CD8^+^ T-cells, respectively^29^.

However, the N protein is more conserved than the S protein, with a sequence conservation of 73.9% across the *Sarbecovirus* and *Merbecovirus* subgenera, compared to 65.8% for the S protein (Fig. 1B). Higher sequence conservation is observed across the surface of the N protein compared to the S protein (Fig. 1C).

### Designing a single antigen vaccine to target *Sarbeco*- and *Merbecoviruses*

We applied Spectravax to design an N protein vaccine antigen. We formulate the objective of Spectravax as designing a protein sequence that protects as many individuals from the target host population as possible against as many pathogens from the target pathogen population as possible. Here, the target host population was the global human population, and the target pathogen population consisted of *β*-CoVs from the *Sarbecovirus* and *Merbecovirus* subgenera.

Spectravax fragments target pathogen protein sequences into short subsequences called k-mers, mimicking antigen processing by the immune system (Fig. 2A). These k-mers are then filtered and scored using two coverage metrics:

1. **Pathogen coverage**: the fraction of target pathogen protein sequences that contain a given k-mer. This metric measures how widely a k-mer is conserved across the pathogen population, indicating its potential to protect against multiple variants.
2. **Host coverage**: the fraction of individuals in the target host population predicted to present at least *n* peptides from a given set of k-mers (default *n* = 1). This metric assesses the likelihood that individuals can present vaccine-derived peptides on their MHC molecules, determined by peptide-MHC binding predictions combined with HLA allele frequencies.

**Fig. 2:**
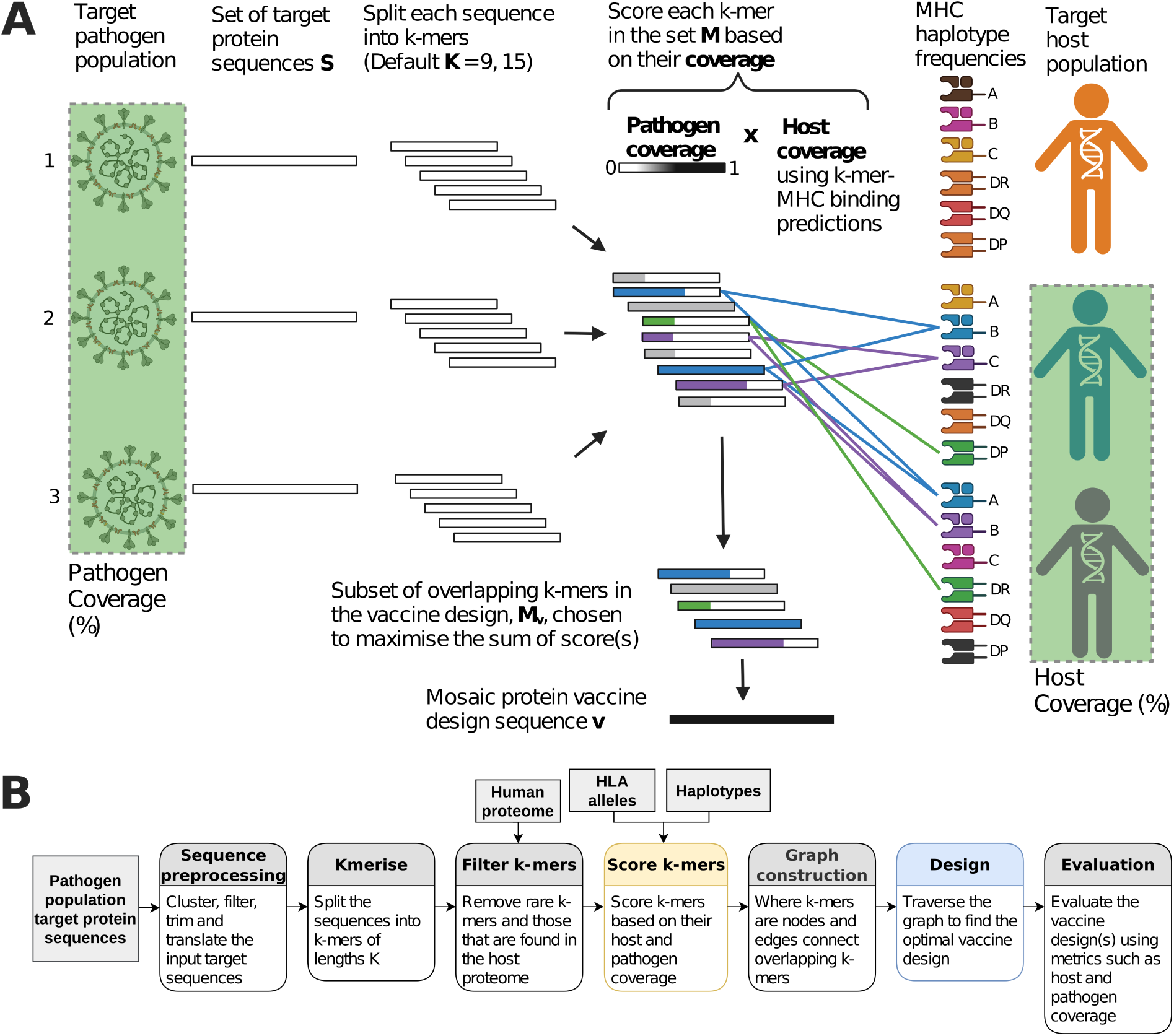
The Spectravax framework. **A)** Overview of the Spectravax vaccine design aiming to maximise coverage of host and pathogen populations. Pathogen coverage is shown as the fraction of pathogen variants (green highlighted viruses) containing peptides in the vaccine design. Host coverage is represented by the fraction of individuals (green highlighted figures) predicted to present vaccine-derived peptides via their HLA alleles. **B)** Spectravax computational workflow: sequence preprocessing, k-mer generation, filtering, scoring, graph construction, vaccine design, and evaluation.

For each k-mer, the host and pathogen coverage scores are multiplied to obtain a total coverage score. The Epigraph algorithm^13^ was reimplemented in Python using the NetworkX library^35^ and used to select the optimal overlapping subset of k-mers to generate the vaccine design (Extended Data Fig. 1).

We chose to design a contiguous protein sequence vaccine rather than a set of peptides or a linker protein vaccine as, by leveraging overlaps, mosaic vaccines can include more epitopes for the same number of amino acids^36;37^, enhancing immunogenicity and avoiding artificial junctions^38^. The Spectravax workflow (Fig. 2B) is further detailed in the Methods section.

### Overview of the Spectravax N vaccine design

The Spectravax N antigen is a protein sequence of 414 amino acids (Fig. 3).

**Fig. 3:**
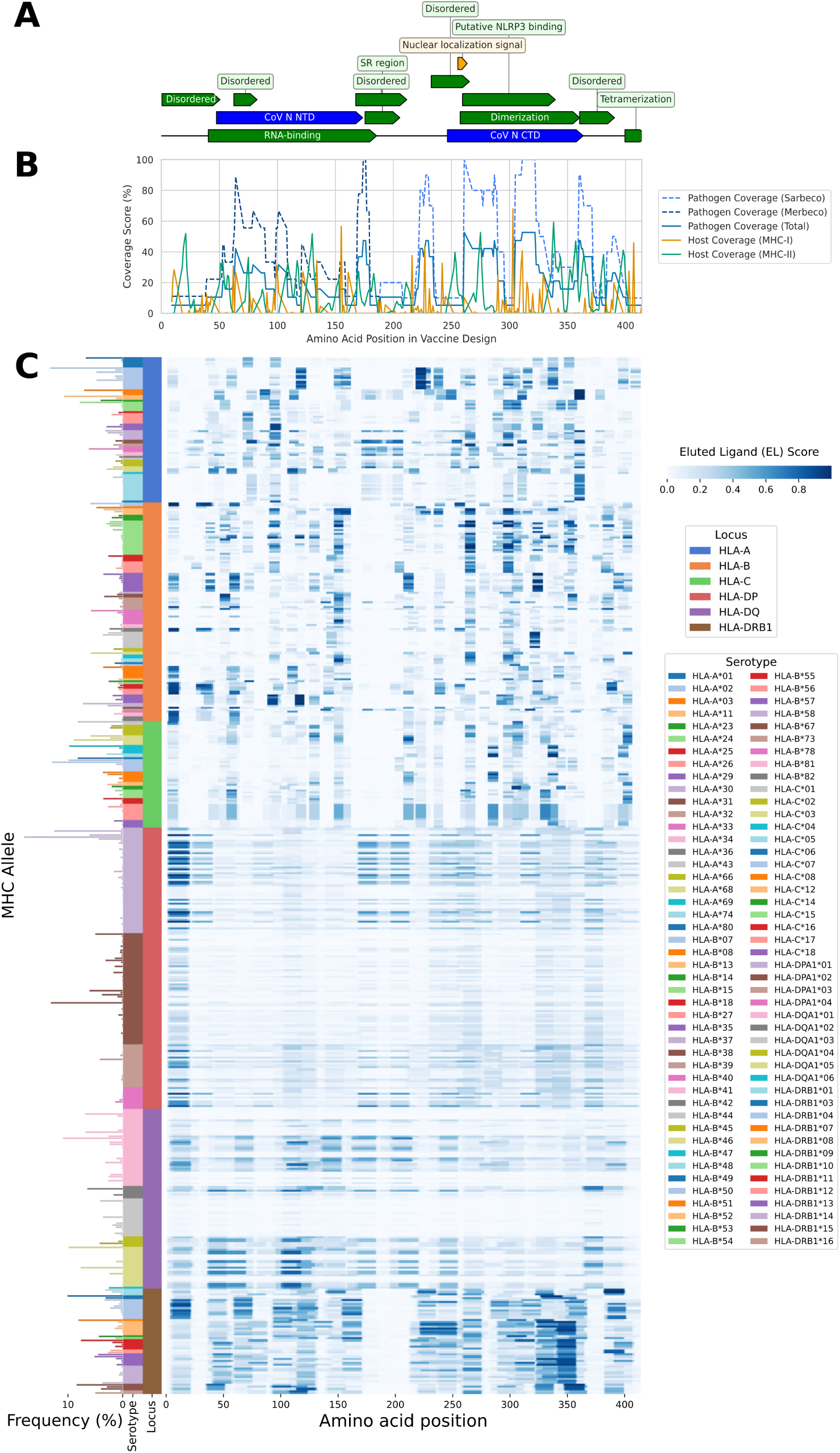
Overview of Spectravax N by amino acid position. **A)** SARS-CoV-2 N protein (UniProt ID: P0DTC9) annotated with domains (blue), regions (green), and motifs (orange). **B)** The Spectravax N antigen, showing k-mer scores for pathogen coverage (blue), *Sarbecovirus* only (light blue), *Merbecovirus* only (dark blue), MHC-I host coverage (orange), and MHC-II host coverage (green). **C)** Peptide-MHC presentation heatmap for Spectravax N. The x-axis represents amino acid position; the y-axis represents HLA alleles, grouped by locus and serotype._8_ The inverted x-axis shows allele frequencies in the global population. The colour intensity indicates the eluted ligand (EL) score, reflecting the likelihood of peptide presentation by MHC molecules.

The pathogen coverage remains high across the designed protein, reflecting its high conservation—all k-mers have non-zero pathogen coverage as they are present in at least one target sequence. In contrast, the host population coverage scores are lower and variable, as only a subset of k-mers are predicted to be presented by MHC class I (MHC-I) or MHC class II (MHC-II) molecules (Fig. 3B).

Intriguingly, the N-terminal domain (NTD) of the antigen design consists entirely of *Merbecovirus* 9-mers, while the C-terminal domain (CTD) contains only of *Sarbecovirus* 9-mers, with a transition at amino acid position 188 (Fig. 3A,B). This separation in the antigen design reflects the distinct nature of the subgenera, which causes each subgenus to have its own region in the antigen design to maximise the coverage of both.

The antigen contains numerous k-mers predicted to be presented by a wide range of MHC alleles (both MHC-I and MHC-II), essential for effective host coverage (Fig. 3C).

The heatmaps reveal ‘hotspots’ where certain regions of the protein are more likely to be presented by MHC molecules. For instance, amino acid positions 5–11 and 262–270 show higher predicted presentation (mean eluted ligand [EL] score over 0.2), while regions like 185–196 and 408–414 show lower presentation (mean EL score below 0.05).

Differences between MHC-I and MHC-II loci are evident in the peptide-MHC heat-maps. MHC-I loci (HLA-A, HLA-B, HLA-C) display more distinct and punctuated patterns, with pronounced variation in EL scores between adjacent k-mers. In contrast, MHC-II loci (HLA-DP, HLA-DQ, HLA-DR) show smoother patterns due to their broader peptide-binding specificity.

Notably, the HLA-DRB1 locus has the highest EL scores, with a mean of 0.23 across all k-mers and alleles—over twice that of other loci (Post-Hoc Dunn test, adjusted p-values *<* 0.001). This may be due to biological factors or biases in the NetMHCIIpan training dataset, as HLA-DRB1 alleles are overrepresented in the training dataset^14^.

Alleles sharing the same serotype exhibit similar peptide presentation patterns due to shared peptide-binding motifs. High predicted presentation scores are observed for serotypes like HLA-DRB1*04, *09, and *10 while others like HLA-DQA1*03 show the lowest scores.

### *In silico* analyses of Spectravax N

We conducted computational analyses to evaluate the predicted coverage of Spectravax N across both host and pathogen populations, and to compare it with wild-type sequences and an existing computational design.

Spectravax N provides high predicted coverage across diverse human populations (Fig. 4A–D). To quantify this coverage, we use the expected number of displayed peptides, denoted as E[#DPs], representing the average number of vaccine-derived peptides predicted to be presented by an individual’s HLA molecules.

**Fig. 4:**
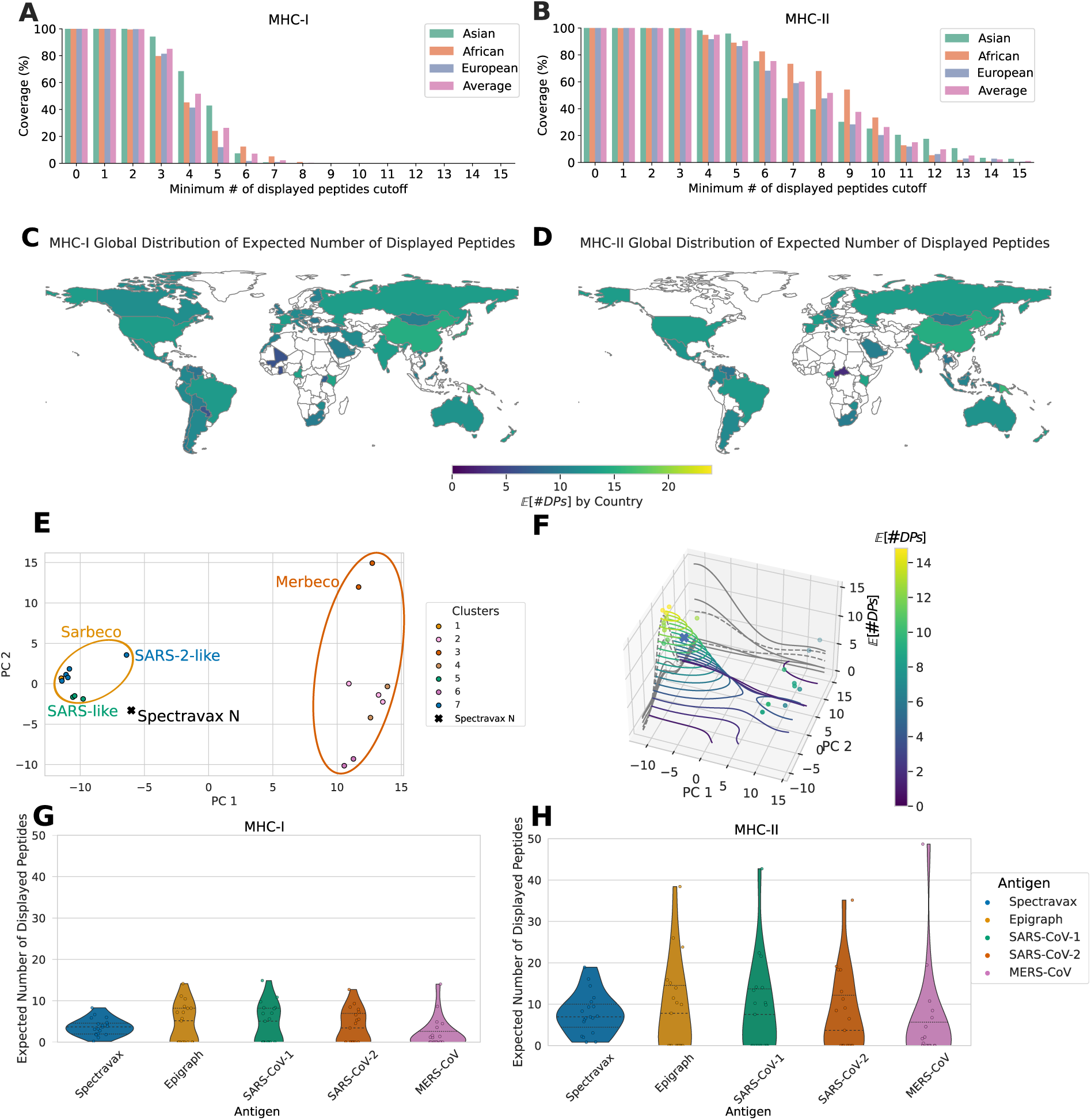
*In silico* analyses of Spectravax N antigen coverage. (**A, B**) Percentage of Asian (green), African (orange), European (blue) individuals, and the combined population (pink) predicted to present at least *n* vaccine-derived peptides for (**A**) MHC-I and (**B**) MHC-II. (**C, D**) Expected number of displayed peptides E[#DPs] across 95 countries for (**C**) MHC-I and (**D**) MHC-II. Countries missing data from the Allele Frequency Net Database (AFND) are coloured white. (**E**) Principal component analysis (PCA) of the N protein sequences from the target pathogens and the Spectravax N antigen in 2D sequence space. Each point represents a target pathogen sequence, coloured by automatically assigned clusters, with the Spectravax N design indicated with a black cross. (**F**) Contour plot of the protein fitness landscape showing the E[#DPs] for each pathogen sequence in the PCA space. (**G, H**) Violin plots of the coverage, showing the E[#DPs] for each antigen against the target sequences for (**G**) MHC-I and (**H**) MHC-II. Colours represent different antigens: Spectravax (blue), Epigraph (yellow), SARS-CoV (green), SARS-CoV-2 (orange), and MERS-CoV (pink).

Firstly, the percentage of individuals with Asian, African, and European ancestries predicted to present at least *n* vaccine-derived peptides for MHC-I and MHC-II was calculated (Figures 4A, 4B). The coverage is consistent across ancestries, with Asian and African populations slightly more likely to present an additional peptide compared to the European population for both MHC classes. Notably, E[#DPs] is consistently higher for MHC-II than for MHC-I across all ancestries. Almost the entire population is predicted to present at least two peptides for MHC-I and at least three peptides for MHC-II, suggesting a robust potential immune response.

Secondly, to assess global coverage, we evaluated E[#DPs] using an independent dataset of allele frequencies from the Allele Frequency Net Database (AFND)^39^ across 95 countries (Figs. 4C and 4D). Despite missing data for some countries, particularly in Africa, the strong predicted responses in highly populous countries like China and India contribute substantially to a high global E[#DPs] of 13.2 when weighted by population size. The rankings of E[#DPs] across ancestries was found to be consistent across datasets, reinforcing the robustness of our findings (Extended Data Table 1).

Thirdly, we evaluated the Spectravax N antigen’s predicted coverage across the target pathogen population, consisting of *β*-CoVs from the *Sarbecovirus* and *Merbecovirus* subgenera, finding that the vaccine design is representative of both subgenera (Fig. 4E, F). This result is further confirmed by the phylogenetic tree and sequence identity matrix (Extended Data Fig. 2).

To visualise the antigen’s placement within the pathogen sequence space, we performed principal component analysis (PCA) on the aligned N protein target sequences. When the distribution of pathogen sequences is presented in 2D space, with each point representing a sequence coloured by automatically assigned clusters, Spectravax N is positioned between the *Sarbecovirus* and *Merbecovirus* clusters (Figure 4E).

To further assess the pathogen coverage of Spectravax N and the target sequences, we calculated the expected number of displayed peptides (E[#DPs]) for each sequence. Figure 4F presents a contour plot mapping the E[#DPs] values onto the 2D sequence space. In this plot, the first two dimensions correspond to the principal components from the PCA, representing sequence similarity, while the contours and colour indicate the E[#DPs] values, analogous to a protein fitness landscape. Spectravax N is located near, but not at, the peak of the fitness landscape due to the clade adjustment process (Extended Data Fig. 3), which balances representation from both subgenera.

Finally, we compared Spectravax N to wild-type N sequences from SARS-CoV, SARS-CoV-2, MERS-CoV, and an antigen designed using the Epigraph method for the same initial set of target sequences (Figs. 4G, H). The Spectravax antigen shows superior breadth, being the only antigen expected to display a non-zero number of peptides against all target sequences. While the Epigraph design and SARS-CoV have a higher E[#DPs] than the Spectravax design, they fail to cover *Merbecoviruses*effectively. Spectravax N, therefore, offers a balanced approach, ensuring coverage across the diverse *β*-CoV population.

### Immunogenicity studies in mice

To test our computational predictions, we experimentally assessed the breadth of the Spectravax N antigen compared to wild-types and an Epigraph-designed antigen using the C57BL/6J mouse model (Fig. 5A). Mice were immunised with pEVAC^40^ plasmid DNA encoding one of the antigens (Spectravax N, Epigraph N, SARS-CoV-2 N or MERS-CoV N) or no antigen (negative control) (Fig. 5B). Mice were immunised three times at four weeks intervals and the study terminated at week twelve, with subsequent sera and splenocytes collection from all the mice groups (Fig. 5A) to characterise the antibody and T-cell responses, respectively. Mice sera were evaluated for IgG antibody binding responses using antigen-specific ELISA (Fig. 5C–E) and mice splenocytes were evaluated for T-cell response using IFN-*γ* ELISpot assays and flow cytometry (Fig. 5F–M).

**Fig. 5:**
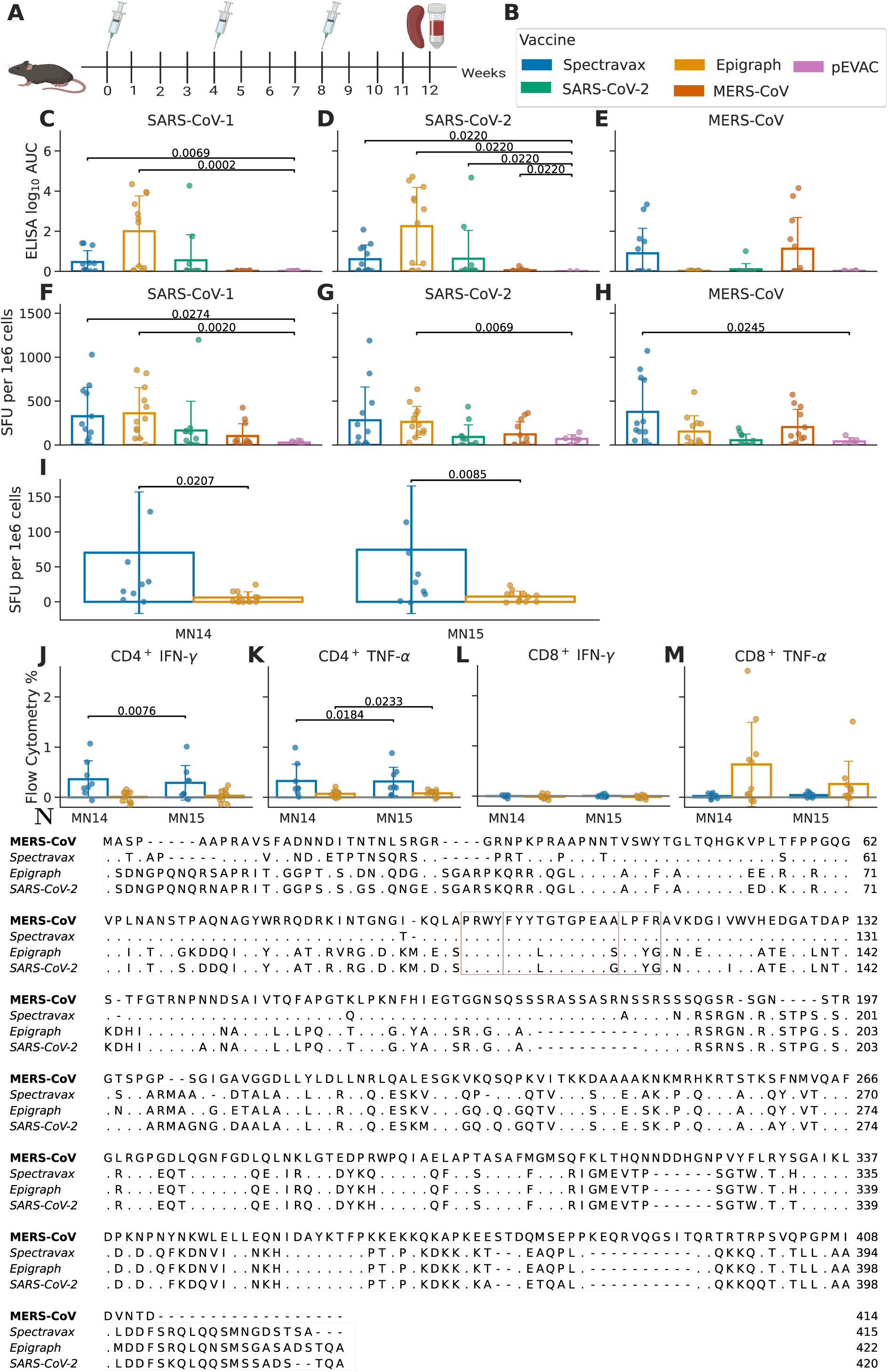
*In vivo* immunogenicity study in mouse model. (**A**) Immunisation schedule. (**B**) Immunisation groups and their colour codes: Spectravax (blue), Epigraph (yellow), SARS-CoV-2 (green), MERS-CoV (orange), and negative control (pink). (**C–E**) ELISA results showing antibody responses against wild-type (**C**) SARS-CoV, (**D**) SARS-CoV-2, and (**E**) MERS-CoV N antigens. (**F–H**) ELISpot results showing T-cell responses measured by IFN-*γ* production upon stimulation with N peptide pools from (**F**) SARS-CoV, (**G**) SARS-CoV-2, and (**H**) MERS-CoV. Statistical significance was determined using a two-sided Mann–Whitney *U* test comparing the Spectravax and Epigraph groups to the negative control. (**I**) ELISpot stimulation with MERS-CoV Nucleocapsid (MN) peptides MN14 and MN15 only (Spectravax: *n* = 10, Epigraph: *n* = 12). (**J–M**) Flow cytometry analysis of the following subpopulations of splenocytes: (**J**) CD4^+^ IFN-*γ*+, (**K**) CD4^+^ TNF-*α*+, (**L**) CD8^+^ IFN-*γ*+, and (**M**) CD8^+^ TNF-*α*+ in response to MN14 and M15 peptide stimulation (Spectravax n=8, Epigraph n=10). P-values are shown only for significant differences (*p <* 0.05). (**N**) Multiple sequence alignment of the antigens generated using ClustalO^41^. The regions corresponding to MN14 and MN15 are boxed in red and blue.

Spectravax N induced antibody responses against SARS-CoV, SARS-CoV-2 and MERS-CoV with antibody titres comparable to the homologous wild-type controls. Epigraph N induced robust and stronger antibody responses against the *Sarbecoviruses* but negligible humoral responses against MERS-CoV, while SARS-CoV-2 and MERS-CoV wild-type antigens generated antibodies only against homologous antigens (Fig. 5C–E). As up to 50% of the mice in every immunised group were non-responders, statistical significance could not be established reliably. The antibody responses elicited in responding mice from the Spectravax and Epigraph groups suggest that some surface-exposed epitopes are preserved in both designs, and the Spectravax design has retained some MERS-CoV B-cell epitopes which are not present on Epigraph (Extended Data Fig. 4).

T-cell responses were initially assessed using ELISpot assays measuring IFN-*γ* production upon stimulation with peptide pools derived from SARS-CoV, SARS-CoV-2, and MERS-CoV N antigens (Fig. 5F–H). Spectravax N induced significantly higher T-cell responses against SARS-CoV and MERS-CoV compared to the negative control (*p <* 0.05) while Epigraph N induced significant T-cell responses against the *Sarbecovirus* antigens (*p <* 0.01), but not against MERS-CoV N.

To identify the peptide(s) that led to differential response to MERS-CoV by the Spectravax and Epigraph antigens, we divided the peptide pools of wild-type MERS-CoV N into smaller peptide pools, with each peptide pool consisting of five to eight peptides. Two mice from each of the Epigraph and Spectravax vaccinated mice groups were selected for the initial ELISpot-based screening (Extended Data Fig. 5A). Only the peptide pool MN11-15 elicited T-cell responses for both antigens, with Spectravax N eliciting a stronger response compared to Epigraph N. Further, the splenocytes from the two same mice were tested for responses against individual peptides which constituted the MN11-MN15 pool (Extended Data Fig. 5B). Responses were observed only for peptides MN14 and MN15, which were computationally predicted to have higher presentation scores (Table 1).

**Table 1:**
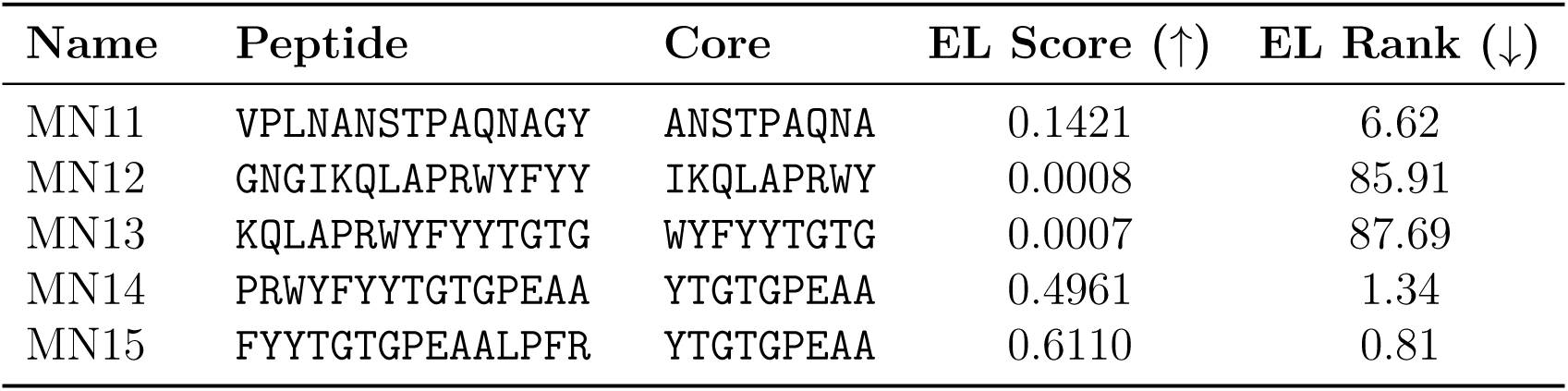
MERS-CoV Nucleocapsid (MN) peptide pool that elicited ELISpot responses. The table lists the peptides, their core sequences, predicted eluted ligand (EL) scores, and percentile ranks. Predictions were made using NetMHCIIpan v4.0^14^ for the C57BL/6J mice MHC-II allele H2-IAb. A higher EL Score (↑) and lower EL Rank (↓) indicates a higher likelihood of presentation.

Mice immunised with either Spectravax or Epigraph antigens were then stimulated with the two identified peptides (Fig. 5I). Significantly higher T-cell responses to peptides MN14 and MN15 were observed for Spectravax-immunised animals compared to Epigraph (*p <* 0.05).

To further delineate which arm of T-cell immunity is stimulated by peptides MN14 and MN15, flow cytometry experiments were performed for mice groups immunised with either Spectravax or Epigraph antigens (Fig. 5J–M). CD4^+^ biased responses were observed for Spectravax N while CD8^+^ biased responses were observed for Epigraph N.

Detailed analyses of peptides MN14 and MN15 show that they have the same core region: ‘YTGTGPEAA’. These peptides have predicted EL scores of 0.5 and 0.61 respectively to mice MHC-II allele H2-IAb and are an exact match between MERS-CoV and Spectravax N, while Epigraph and SARS-CoV-2 antigens have mismatches at two positions in the core region and four positions over the length of the region corresponding to the peptides (Fig. 5N). Furthermore, these epitopes may be important for the target human population as MN14 and MN15 have an expected MHC-II host coverage (Extended Data Section 1) of 30.8% and 10.3% respectively.

These results demonstrate that even small differences in epitope sequences can substantially impact T-cell responses. This emphasises the importance of computational methods that can enrich antigen sequences with optimal T-cell epitopes, particularly for generating broad-spectrum vaccines capable of protecting against multiple target pathogens.

Computational predictions made prior to experimental validation showed a strong correlation with the experimental results, particularly for the ELISA responses: MHC-I *R_S_* = 0.86, MHC-II *R_S_* = 0.89 (Extended Data Fig. 6). This correlation suggests that computational predictions can effectively guide vaccine design before time- and resource-intensive experimental validation.

## Discussion

Broad-spectrum vaccines are crucial for combating antigenically diverse pathogens with pandemic potential. Computational methods for designing single-antigen broad-spectrum vaccines offer a generalisable approach that, in theory, can be applied to any pathogen. In this work, we introduced Spectravax, a computational framework to design broad-spectrum vaccines that accounts for the genetic diversity present in both pathogen and host populations. The Spectravax framework has a strong theoretical foundation and is intuitive, as the scores correspond to real-world quantities such as population coverage and the number of displayed peptides. This makes the results interpretable and circumvents the need for arbitrary weights used in other methods^38^. Furthermore, the underlying framework is generalisable so it can be easily extended by other researchers. To demonstrate its utility, we applied Spectravax to design a N vaccine antigen to target *Sarbeco*- and *Merbecoviruses*.

One of the main advantages of Spectravax compared to existing methods is its use of the host coverage score, which maximises T-cell epitope presentation across a target population. A larger independent dataset of allele frequencies from AFND was used to validate the consistency of the host coverage estimates. Rarer HLAs, which tend to be less well studied and thus have less confident binding predictions, receive less weight in the host coverage score, thus increasing the global coverage. Antigens designed using Spectravax, such as the N antigen, were predicted to display peptides across a wide range of HLA alleles, leading to effective host coverage. This was demonstrated for populations from three ancestries and 95 countries. Incorporating the host coverage score consistently improved the expected number of displayed peptides across multiple antigens, although pathogen coverage was found to be more important (Extended Data Fig. 7). Another major improvement over existing methods is the use of clade equalisation, which helps mitigate biases towards clades with higher number of characterised members, such as the *Sarbecovirus* subgenus in the case of *β*-CoVs. Interestingly, this process resulted in a chimera-like N antigen combining sequences from the *Merbecovirus* and *Sarbecovirus* clades at the N and C termini, respectively. This design was found to be representative of both subgenera and was the only evaluated antigen able to display peptides against all target sequences. This broad coverage was experimentally validated in mice using representatives of SARS-CoV, SARS-CoV-2, and MERS-CoV N proteins. Although the Epigraph and SARS-CoV-2 antigens induced stronger SARS-CoV and SARS-CoV-2 responses, the response against MERS-CoV was not statistically significant. Furthermore, we were able to identify the peptides responsible for the difference in response to MERS-CoV. On evaluating the protein fitness landscape (Fig. 4F) illustrates the tradeoff between potency and breadth, an observation made with some of our earlier vaccine designs as well^10^.

Limitations of the Spectravax method primarily stem from the underlying sequencing and immunological data used. The dbMHC frequency data used to estimate host coverage covers only 15 geographic regions and may not fully reflect the global human population^42^. Mass spectrometry data is limited to the peptides that are eluted, which may not represent the full spectrum of peptides displayed *in vivo*, and may be biased towards more common HLAs^14^. Pathogen sequence databases are known to have biases such as over-representation of certain viral families, geographic regions, hosts, or time periods^43^. The method also requires substantial computational resources for peptide-MHC binding predictions. Fortunately, the data limitations of Spectravax may diminish over time as more sequencing and immunological data become available.

In terms of future work, Spectravax can be applied to other pathogens and host species, as evidenced by its application to seven different antigens from two diverse viral families (Extended Data Table. 2). The pathogen coverage score could be improved by incorporating pathogen prevalence^13^ or spillover risk^44^, similar to clade equalisation. The host coverage score could incorporate predictions for T-cell activation^45;46;47^, in addition to peptide presentation. Although the N antigen demonstrated significant antibody responses, further computational optimisation could be done to enhance these, either by integrating methods for linear B-cell epitope prediction^18^ or by using generative deep learning models to design structurally conserved antigens while optimising for both host and pathogen coverage^17^.

In conclusion, Spectravax N represents the first computationally designed antigen to elicit immune responses against SARS-CoV, SARS-CoV-2, and MERS-CoV, highlighting its potential as a pan-*β*-CoV vaccine candidate. Apart from Spectravax N, we are aware of only one other single-antigen vaccine, consisting of a SARS-CoV-2 S protein with a MERS-CoV RBD, that has shown to protect against SARS-CoV, SARS-CoV-2 Omicron variant and MERS-CoV challenge in mice^48^. The Spectravax method provides a substantial advancement in the rational design of broad-spectrum vaccines that can be applied to other pathogens and host species. The insights gained from this work regarding host and pathogen coverage offer valuable lessons for the broader field of vaccine development and can contribute to improved pandemic preparedness by supporting the development of vaccines that can protect against emerging infectious diseases.

## Methods

### Spectravax framework

Spectravax is a computational method to design broad-spectrum vaccines by maximising the coverage of both pathogen and host populations, accounting for genetic diversity across both.

The vaccine design problem is formulated as follows^13;36^:

Let *S* = {*s*_1_*, s*_2_*, . . ., s_N_* } represent the set of protein sequences from the target pathogen population. Each protein sequence *s_i_* is split into overlapping k-mers of different lengths from a set *K* = {*k*_1_*, k*_2_*, . . ., k*_|_*_K_*_|_}. The union of all k-mers in the pathogen sequences is denoted as *M* (*S*). The objective is to design a contiguous protein sequence *v* composed of k-mers from *M* (*S*) that maximises the total coverage score, subject to the constraint shown below. The constraint ensures that any two consecutive *k*-mers in the vaccine design share an overlap of *k* − 1 amino acids, where *k* is the shorter *k*-mer length. This ensures contiguity of the amino acid sequence while accommodating varying *k*-mer lengths.

The optimisation problem is expressed as:

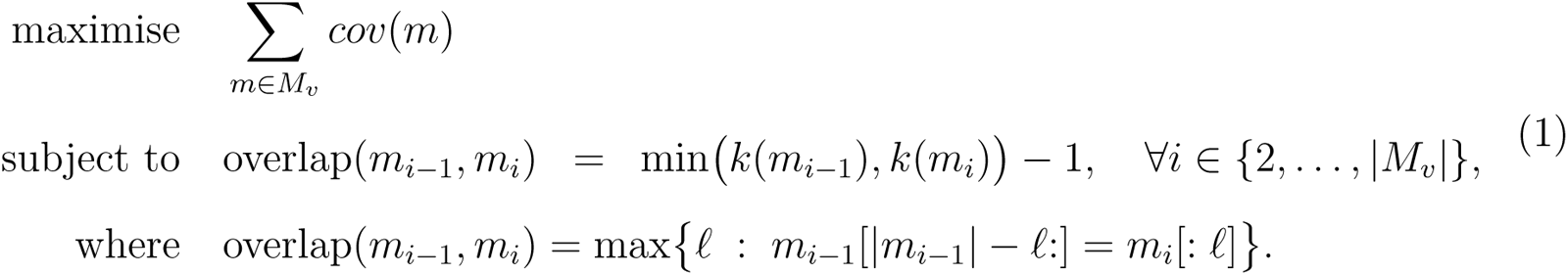

Where *M_v_* is the set of k-mers in the vaccine design *v*, and *cov*(*m*) is the total coverage score for k-mer *m*, which is calculated as the product of the pathogen and host coverage scores:

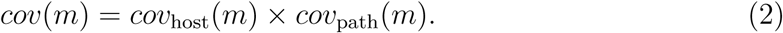

### Pathogen population coverage

The pathogen coverage score (*cov*_path_) is calculated as the fraction of target pathogen sequences containing the k-mer^13^:

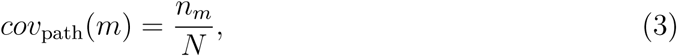

where *n_m_* is the number of target sequences that contain the k-mer *m*, and *N* is the total number of target sequences. This gives a score between 0 and 1, where 1 indicates that the k-mer is fully conserved across the target pathogen population. This is also referred to as conservation in the wider literature^36^, however, we use the term pathogen coverage to emphasise similarities with the host coverage score.

### Host population coverage

The host coverage score (*cov*_host_) is computed as the weighted sum of the MHC-I (*cov*_mhc1_) and MHC-II (*cov*_mhc2_) coverage scores:

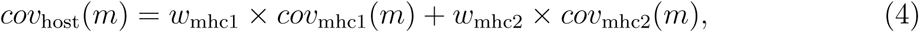

where *w*_mhc1_ and *w*_mhc2_ are the respective weights for MHC-I and MHC-II scores, set to 1 unless specified otherwise. This term is often simply referred to as population coverage in the literature^15^, however, we use the term host coverage to differentiate it from the pathogen coverage score.

The MHC-I and MHC-II coverage scores are calculated using the EvalVax-Robust algorithm^15^, which accounts for linkage disequilibrium between HLA alleles by using haplotype frequencies. The method combines peptide-MHC binding predictions with host HLA allele frequencies to estimate the proportion of the host population that can present each k-mer. Host HLA allele frequencies were collected from the dbMHC database, covering three main ancestries (Asian, African, and European)^42^. For validation across broader populations, allele frequencies from the Allele Frequency Net Database (AFND) covering 95 countries were also used^39;49^. However, unlike EvalVax-Robust^15^, we use the eluted ligand (EL) binding score as the binding criteria, as described below. This enabled the use of the expected host coverage score, which increased the expected number of displayed peptides (Extended Data Section 1).

### Peptide-MHC binding predictions

Binding predictions were performed using NetMHCpan v4.1 for MHC-I and NetMHCIIpan v4.0 for MHC-II^14^. The eluted ligand (EL) score was selected as the binding criteria, as it was shown to be discriminative in identifying binders for SARS-CoV-2 peptides and well calibrated (Extended Data Fig. 8)^15^. Peptide-HLA binding predictions were made for all relevant HLA alleles from the dbMHC database^42^. Due to the computational intensity of these predictions, calculations were parallelised over both k-mers and HLA alleles using the redun workflow management system^50^.

### Sequence data and processing

Protein sequences of the nucleocapsid (N) protein from the *Sarbecovirus* (taxid: 2509511) and *Merbecovirus* (taxid: 2509494) subgenera were retrieved from the NCBI Virus database^30^. Sequences were filtered to include only entries with the relevant protein names: “nucleocapsid phosphoprotein”, “nucleocapsid protein”, “nucleoprotein”, and “nucleocapsid”. Sequences with incomplete or ambiguous amino acids were removed. To reduce redundancy and ensure a representative set of sequences, nucleotide sequences were clustered using CD-HIT^51^ at a threshold of 99% identity, translated to amino acids and duplicate protein sequences were removed. After preprocessing, 84 target sequences remained for vaccine design.

### Antigen design

We applied Spectravax to design a vaccine antigen targeting the *Betacoronavirus* (*β*- CoV) nucleocapsid (N) protein. For the N antigen, clade equalisation was employed to ensure balanced representation of the *Sarbecovirus* and *Merbecovirus* subgenera, fixing the number of clusters to seven. This ensures the final design incorporates conserved elements from both subgenera. The sequences were further clustered at 95% within the Spectravax before clade equalisation, resulting in a final set of 19 target sequences. The MHC-I weight was set to double that of MHC-II, in an attempt to focus on eliciting a CD8^+^ T-cell response. All other parameters were set to their default values. For all other vaccine designs (Extended Data Fig. 9), the default parameters were used (Extended Data Table 3).

### Computational analyses

The expected number of displayed peptides (E[#DPs]) is computed by summing over all thresholds of *n* displayed peptides, weighted by the corresponding coverage at each threshold^15^:

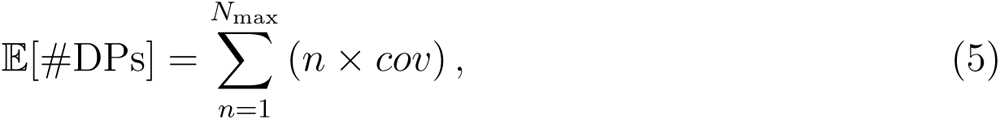

where *cov* is the coverage, i.e. the fraction of the population that is predicted to display exactly *n* peptides, and *N*_max_ is the maximum possible number of peptides.

Principal component analysis (PCA) was performed to visualise the diversity of pathogen sequences and to position the vaccine designs within sequence space. A multiple sequence alignment (MSA) was first generated using MAFFT^31^. The MSA is then converted to a numerical matrix where the amino acids at each position are ranked by their frequency in the MSA. PCA was applied to this feature matrix to reduce dimensionality, using the Python package scikit-learn^52^ and the first two principal components were plotted.

### Plasmid DNA preparation

Genes encoding the antigens were commercially synthesised and cloned into the pEVAC vector (GeneArt, Thermo Fisher Scientific). The N sequences were codon-optimised for expression in mammalian cells using the GeneOptimizer algorithm^53^. Plasmid DNA used for vaccination were outgrown and isolated using EndoFree Plasmid Mega Kit (Qiagen) following the manufacturer’s instructions. The extracted plasmids were then quantified using UV spectrophotometry with a NanoDrop device (Thermo Scientific)^53^. The presence of endotoxin was measured using the Pierce LAL Chromogenic Endotoxin Quantitation Kit (Thermo Scientific) and was found to be below the detection limit of 20 EU/ml. The sequences were confirmed using Sanger sequencing (Source BioScience).

### Immunisation of mice

Four groups of twelve 7–10-week-old female C57BL/6J mice, plus an additional six mice for the control group, were obtained from Charles River Laboratories (Margate, U.K). All animal experiments were conducted in compliance with institutional guidelines and approved by the Animal Welfare and Ethical Review Board covering University Biomedical Services, University of Cambridge. Mice were immunised subcutaneously in the rear flank with a 40 *µ*g dose of plasmid DNA diluted in phosphate-buffered saline (PBS). Immunisations were administered on days 0, 28, and 56, following the schedule shown in Fig. 5A. Terminal blood collection was performed by cardiac puncture under non-recovery isofluorane anaesthesia on day 84, and spleens were harvested using aseptic technique for immunological assays.

### Peptide design and synthesis

Peptide sequence predictions were generated using NetMHCpan and NetMHCIIpan software without a length restriction setting^14;54^. For predicted peptide pools, the library of peptides was filtered for binding values, number of alleles being targeted, and the number of times amino acid was predicted to be a part of the peptide. Once such assembly of peptides was selected, the predicted peptides were aligned to the reference sequence and a peptide library of 15-mer with overlap of 11-mer was generated.

For overlapping peptide pools, entire length of the protein reference sequence was divided into 15-mer peptides with 11-mer overlap. All peptides were synthesized by GenScript and reconstituted in 4% DMSO in PBS.

### Isolation of murine splenocytes

Murine splenocytes were isolated from dissected spleens by mashing spleens on MACS SmartStrainers (Miltenyi Biotec) and layering them on the Histopaque 1083 reagent (SigmaAldrich). Puffy coat containing immune cells was collected from the phase separation region and washed in R10 medium (RPMI medium (Gibco) supplemented with 10% Fetal bovine serum (FBS) (Gibco)). Cell pellets were resuspended and frozen in FBS supplemented with 10% DMSO Hybri-Max (SigmaAldrich).

### ELISpot assay

ELISpot assays with frozen splenocytes were performed according to the manufacturer’s protocol using the mouse IFN-*γ* Single-Color ELISpot kit (Immunospot). Frozen splenocytes were thawed, washed once with R10 medium and resuspended in CTL medium supplemented with L-glutamine at the desired cell density (in the range of 2-5×106 cells/mL). Custom made predicted peptides for full-length SARS-CoV-2 N (NCBI Reference Sequence: YP_009724397.2), SARS-CoV N (NCBI Reference Sequence: YP_009825051.1) and MERS-CoV N (NCBI Reference Sequence: YP_009047204.1) proteins were used. For the stimulation, 100 *µ*L of peptide solution in CTL at 4 *µ*g/peptide/mL were mixed with 100 *µ*L of cell suspension in the 96-well ELISpot plate and incubated for 24h in 37 oC in humidified incubator with 5% CO2. As a positive control, eBioscience Cell stimulation cocktail (ThermoFisher Scientific) was used at 1x dilution. CTL medium was used as a non-stimulation control. The plates were developed as per manufacturer’s protocol, dried, scanned and analysed on ELISpot reader S6 Ultra M2 (ImmunoSpot).

To identify immunodominant peptides for the MERS-CoV N protein, 8 smaller peptide pools were prepared and two samples from each Spectravax and Epigraph vaccinated groups of mice were assayed. After identification of the two specific immunogenic peptides (MN14 and MN15 from the original MERS-CoV N peptide pool), there were used at 2 *µ*g/mL for the final concentration on the entire available cohort of responding animals from Spectravax and Epigraph groups. All samples were assayed in at least technical duplicates.

### Flow cytometry cytokine assay

Under sterile conditions murine splenocytes were placed into RPMI (10% FBS, 1% P/S 50 *µ*M beta-mercaptoethanol) media in a 96 well U-bottomed plate at a concentration of 1.5×106/ml, 200 *µ*l per well. The cells were incubated either with Protein Transport Inhibitor (Thermofisher 00-4980-03) + 1 *µ*g/ml SARS-CoV, SARS-CoV-2 or MERS-CoV pooled peptides. PTI and CSC were used as a negative and positive control respectively. Cells were incubated at 37°C for 6 hours.

Cells were washed with FACS buffer (PBS, 0.5% BSA, 0.01% NaN3) three times, blocked with FACS buffer with 2% rat serum for 15 minutes at 4°C, then stained with Fixable Viability Dye eFluor™ 780 (ThermoFisher 65-0865-14), CD4 Monoclonal Antibody (RM4-5) APC (ThermoFisher 17-0042-83), CD3e Monoclonal Antibody (145-2C11) PE-Cyanine5.5 (ThermoFisher 35-0031-82) and CD8a Monoclonal Antibody (53-6.7) PE-Cyanine7 (ThermoFisher 25-0081-82). Cells were incubated for 30 minutes at 4°C in the dark, washed three times with FACS buffer, then resuspended in CytoFix/CytoPerm (BD Biosciences 554714) and left at room temperature (RT) in the dark for 15 minutes. Cells were then washed with perm/wash buffer 3 times, stained for intracellular cytokines with IFN-*γ* Monoclonal Antibody (XMG1.2) Alexa Fluor™ 488 (ThermoFisher 53-7311- 82) and TNF-*α* Monoclonal Antibody (MP6-XT22), PE-eFluor™ 610 (ThermoFisher 61-7321-82) at RT in the dark for 30 minutes. Washed with perm/wash buffer 3 times, and finally resuspended in FACS buffer for analysis on an Attune NxT Flow Cytometer equipped with 3 lasers (ThermoFisher Scientific). A BD LSRFortessaTM Cell Analyser with 5 lasers (BD Biosciences) was used when analysing the individual reactions to MERS-CoV pools 1-8 at 2 *µ*g/ml each, and MERS peptides MN14 and MN15 at 2 *µ*g/ml each in a separate experiment. UltraComp eBeads™ Compensation Beads were used for single colour compensation controls. Analysis was performed using FlowJo 10.9.0 (BD Biosciences).

### ELISA assay

Serum IgG antibody binding responses were assessed by ELISA against recombinant full length N proteins from SARS-CoV (ACROBiosystems, NUN-S5229), SARS-CoV-2 (kind gift from Dr. Leo C. James), and MERS-CoV (CD Creative Diagnostics, DAG-H10295). Nunc MaxiSorp flat-bottom plates were coated with 1 *µ*g/ml of recombinant N protein in PBS (-/-) and incubated overnight at 4°C. Plates were blocked with 3% milk in PBST (PBS with 0.1% Tween 20) for 1 hour at 300 RPM. 50 *µ*l of serum samples diluted in PBST with non-fat milk was added and incubated for 2 hours at room temperature. Plates were washed three times with PBST, and HRP-conjugated goat anti-mouse IgG (H+L) secondary antibody (Jackson Immunoresearch, 115-035-003; 1:3000 dilution) was added to each well. After incubation for 1 hour at room temperature, plates were washed three times with PBST, and 50 *µ*l of TMB substrate was added to each well. The reaction was stopped after three minutes by adding 50 *µ*l of 1 M H_2_SO_4_, and the absorbance was read at 450 nm using a BioTek 800TS microplate reader.

### Statistical analysis

Statistical significance was determined using a two-sided Mann-Whitney *U* test for comparisons between groups using the SciPy library in Python. All plots were generated using the Matplotlib library in Python.

## Acknowledgements

P.P. was supported by a fellowship provided by Open Philanthropy. We thank Dr. Leo C. James for the provision of purified recombinant SARS-CoV-2 nucleoprotein.

## Author Contributions

P.P. and S.V. conceptualised the study, developed the Spectravax method, designed the antigens and generated the figures. P.P., S.V., S.F., G.W.C. and J.L.H. planned the experiments. J.L.H. acquired funding. S.V., S.F., G.W.C. and J.L.H. supervised the project. C.G., S.P. and P.P. performed DNA purification and preparation. L.O. and G.W.C. performed the animal work. S.P., C.G., J.H., L.W. and S.F. performed the *in vitro* assays. S.F. and J.H. analysed the assay data. P.P. wrote the original draft of the manuscript. S.V., G.W.C., S.F. and J.L.H. reviewed and edited the manuscript. All authors provided feedback on the paper.

## Extended Data

**Extended Data Fig. 1:**
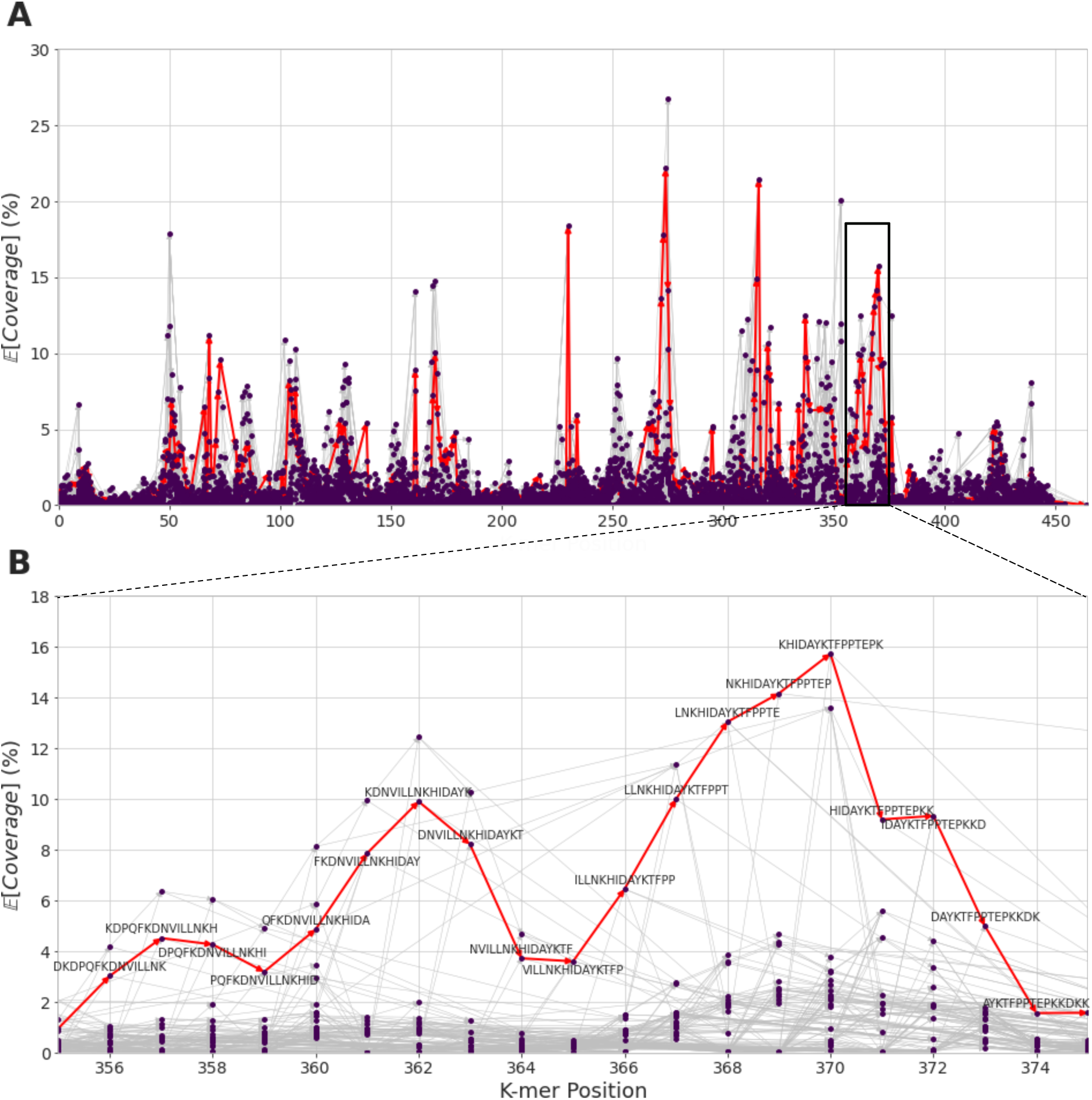
K-mer graph for the Spectravax N antigen design. (**A**) K-mer graph across the entire sequence. The section highlighted in black rectangle corresponds to the region shown in subplot B. (**B**) K-mer graph for amino acids 355-375. The nodes of the graph are the dots that represent all unique k-mers. The edges are the thin lines that connect k-mers whose sequence overlap by *k* − 1 amino acids. The red lines represent the optimal path through the graph, which corresponds to the vaccine design. The x-axis represents the position in the sequence, the y-axis represents the expected coverage score E[*cov*] (Extended Data Section 1). The position (x-axis) is approximate as a given k-mer can appear at different positions in different sequences. Therefore, the k-mer position was measured by the first instance found in a multiple sequence alignment of the target sequences generated using MAFFT^31^.

**Extended Data Table 1:** Comparison of the expected number of displayed peptides (E[#DPs]) for different antigen designs and ancestries/continents using the dbMHC and AFND databases. The Spectravax antigen was designed and evaluated using haplotype frequencies originally from the dbMHC database. To validate the robustness of the results, an independent dataset of allele frequencies from the Allele Frequency Net Database (AFND) was used to evaluate the E[#DPs] for 95 countries with available data. This table shows a comparison of the E[#DPs] for the antigen design and ancestries/continents using the two databases. The results computed using the two databases are similar, with consistent rankings of the E[#DPs] for all ancestries and antigens, supporting the reliability of the results obtained using the dbMHC database. However, the E[#DPs] computed using AFND are consistently higher than those computed using dbMHC. This discrepancy may be due to the different methods used, as the AFND calculations are based on diploid frequencies, whereas dbMHC calculations are based on haplotype frequencies.

| Antigen | Ancestry | $\mathbb{E}[\#DPs] \uparrow$ | |
| --- | --- | --- | --- |
|  |  | dbMHC | AFND |
| Spectravax N | Asian | 11.9 | 14.4 |
|  | African | 11.8 | 13.6 |
|  | European | 10.6 | 13.4 |

**Extended Data Fig. 2:**
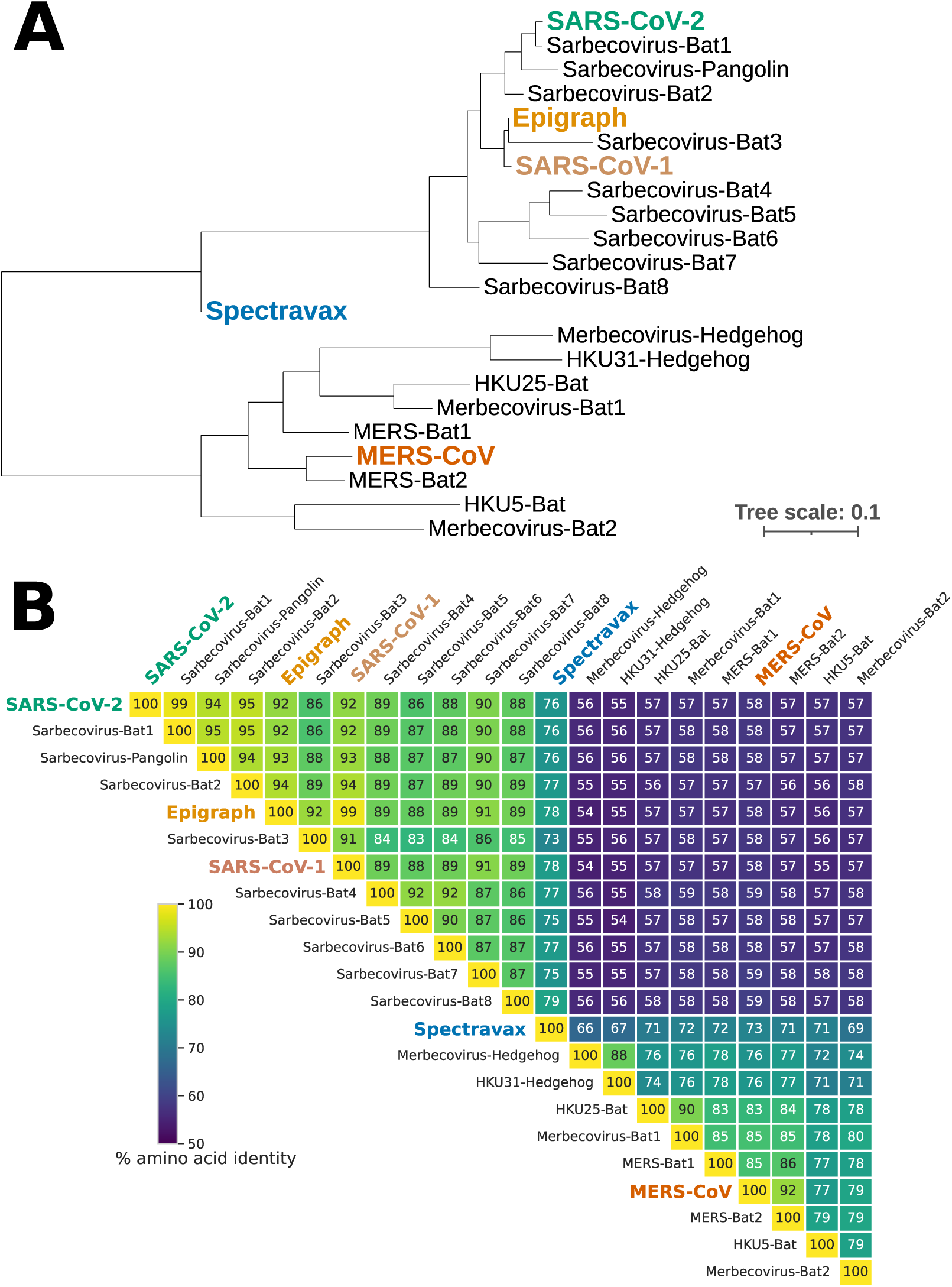
Phylogenetic tree and sequence identity matrix of *Sarbecovirus* and *Merbecovirus* Nucleoprotein sequences. (**A**) Phylogenetic tree produced by multiple sequence alignment (MAFFT^31^) and constructed in IQ-TREE 2^55^ using the LG model^56^, then visualised with iTOL^57^ with midpoint root. (**B**) Sequence identity matrix generated by pairwise alignment using Biopython’s globalxx method^58^. The Nucleoprotein sequences are those used as the target of vaccine design and supplemented with: Spectravax design (blue), Epigraph design (yellow), SARS-CoV-1 (brown), SARS-CoV-2 (green), and MERS-CoV (orange). The GenBank protein IDs for the target proteins are as follows: QSQ01658.1 (Sarbecovirus-Bat1), QVT76614.1 (Sarbecovirus-Pangolin), QZX47293.1 (Sarbecovirus-Bat2), QZX47283.1 (Sarbecovirus-Bat3), YP_003858591.1 (Sarbecovirus-Bat4), QVN46576.1 (Sarbecovirus-Bat5), QTJ30142.1 (Sarbecovirus-Bat6), BCG66635.1 (Sarbecovirus-Bat7), UFP05031.1 (Sarbecovirus-Bat8), YP_009513018.1 (Merbecovirus-Hedgehog), UMO75626.1 (HKU31-Hedgehog), ASL68949.1 (HKU25-Bat), AH2Y461344.1 (Merbecovirus-Bat1), AUM60021.1 (MERS-Bat1), ATQ39393.1 (MERS-Bat2), YP_001039969.1 (HKU5-Bat), ABG47058.1 (Merbecovirus-Bat2).

**Extended Data Fig. 3:**
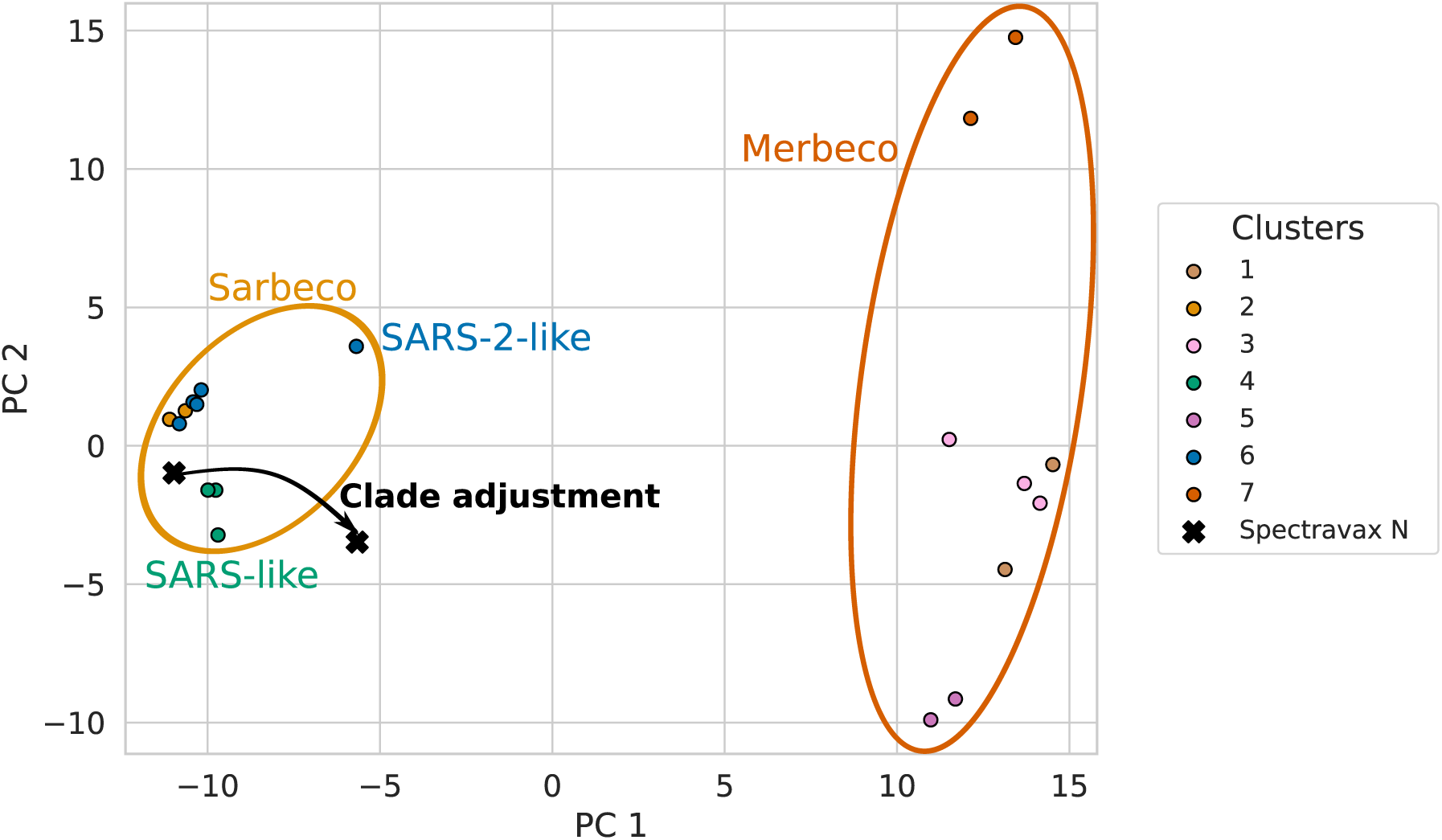
Spectravax clade adjustment. The effect of clade adjustment on the Spectravax N design is shown using principal component analysis (PCA). The x- and y-axes represent the first two principal components of the ranked multiple sequence alignment (MSA) of the *Sarbecovirus* and *Merbecovirus* N sequences. The clade adjustment process (shown as a black arrow) shifts the vaccine design closer towards the centre of the sequence space, enhancing its breadth across the target pathogen population. Clade adjustment therefore addresses bias towards overrepresented clades (e.g., *Sarbecovirus*) by adjusting coverage scores to give higher weight to k-mers from under-represented clades (e.g., *Merbecovirus*).

**Extended Data Fig. 4:**
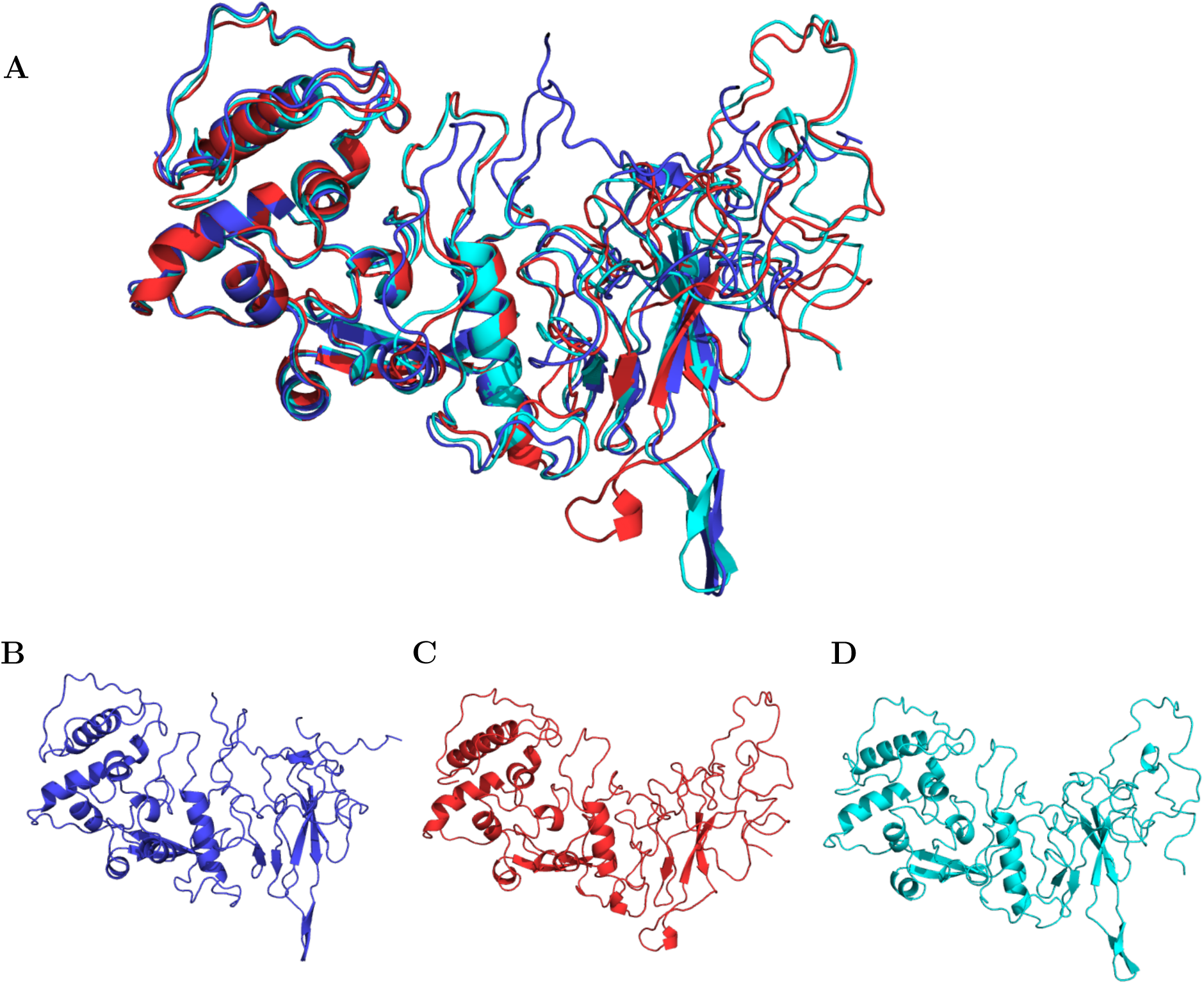
AlphaFold-predicted structures of Spectravax, Epigraph, and SARS-CoV-2 Nucleoproteins. (**A**) Overlay of all structural models: (**B**) Spectravax N (blue), (**C**) SARS-CoV-2 N (red, PDB ID: 8FD5^59^), (**D**) Epigraph N (cyan). Spectravax and Epigraph N structures were predicted with ColabFold^60^ using AlphaFold2^61^ and the SARS-CoV-2 N template: 8FD5^59^. Default parameters were used and the best model was selected based on pTM-score.

**Extended Data Fig. 5:**
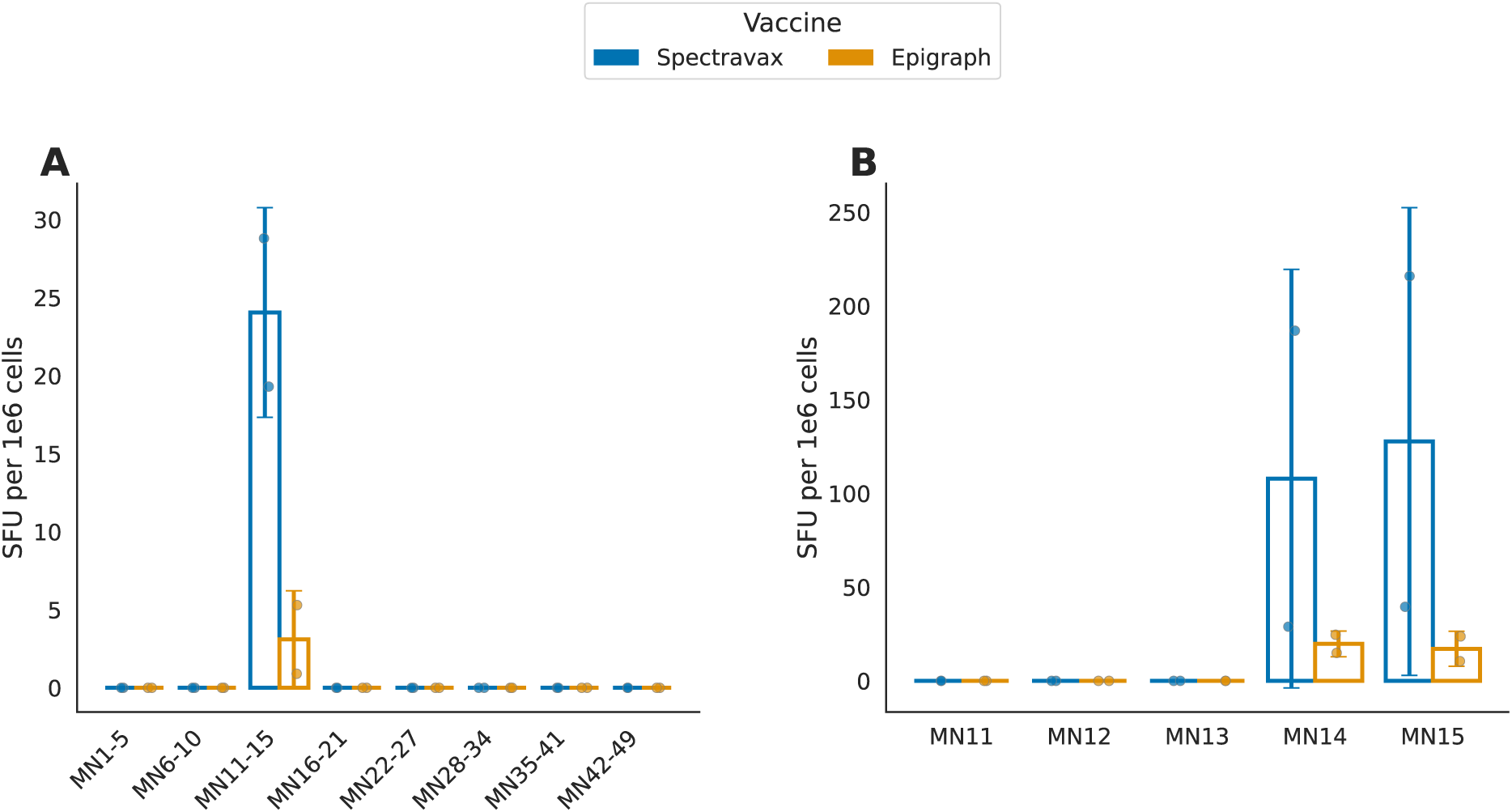
Identification of the immunodominant peptides using ELISpot. (**A**) Stimulation with MERS-CoV Nucleocapsid (MN) peptide pools. (**B**) Stimulation with individual peptides from pool MN11-15 that showed the highest response. Two antigens were evaluated: Spectravax (blue) and Epigraph (yellow, n=2).

### 1 Expected host coverage score

As the EL score is essentially a measure of the likelihood that a peptide is naturally processed and presented on MHC molecules, it can be incorporated directly as a probability in the calculation of host coverage. Unlike the binary hit-based approach used by EvalVax-Robust^15^, where peptides are filtered based on a strict binding affinity threshold, this probabilistic approach uses the EL scores to derive *P_p,a_*, the probability that allele *a* presents peptide *p*. For each allele *a*, let:

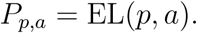

Given a haplotype *h* = *A_i_B_j_C_k_*, the probability that peptide *p* is displayed is:

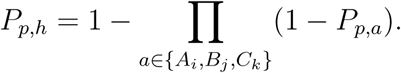

Rather than applying a threshold to determine whether a haplotype is covered by at least one peptide, the probabilities *P_p,h_* are summed over all peptides in *O*:

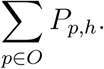

This summation provides the expected number of displayed peptides per haplotype, reflecting the average count of peptides likely to be presented rather than the probability that at least one peptide is presented. When weighting by haplotype frequencies and summing over all haplotypes, we obtain:

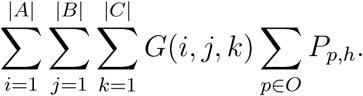

This value represents the expected number of displayed peptides per individual in the population.

If we focus on a single peptide *p*, then:

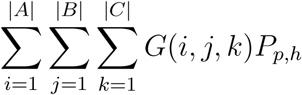

directly yields the expected fraction of the population that displays that peptide.

As shown in Extended Data Fig. 10, using the expected host coverage score (E[Coverage]), rather than the coverage score, increased the expected number of displayed peptides by 0.6 (10.2%) for MHC-I and by 1.1 (7.8%) for MHC-II.

This indicates that expected host coverage provides a more sensitive measure for evaluating individual k-mers. Under the strict thresholding from OptiVax, many peptides have zero 1-times host coverage. In contrast, the expected host coverage is typically non-zero, enabling improved discrimination among candidate peptides. While the expected host coverage was not used to design Spectravax N, as the vaccine designs were ordered before the development of this method, these findings suggest that future Spectravax-based vaccines could show even more promising results.

**Extended Data Fig. 6:**
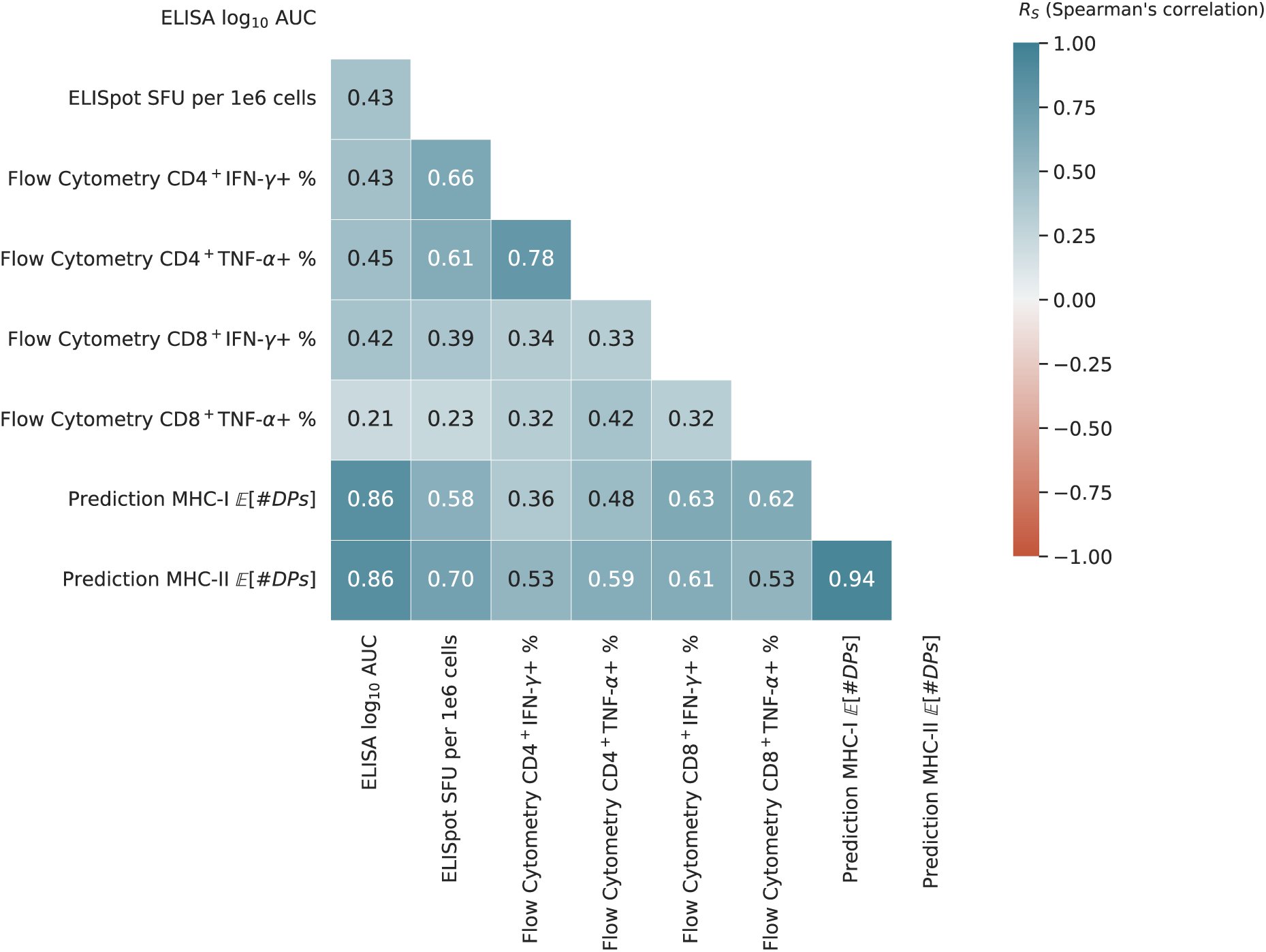
Correlation matrix of experimental results and computational predictions in C57BL/6J mice. The Spearman correlation coefficient (*R_S_*) is shown for each pair of methods. The colour of each cell indicates the strength of the correlation, with blue representing positive correlations and red representing negative correlations. Only positive correlations were observed between the different assays, indicating general agreement in the strength of immune responses measured by different methods. ELISpot responses correlated more strongly with CD4^+^ T-cell responses (*R_S_ >* 0.6) than with CD8^+^ T-cell responses (*R_S_ <* 0.4), suggesting that CD4^+^ cells were the main drivers of the T-cell responses. The experimental results also showed a strong correlation with the computational predictions. In particular, the computational predictions correlated strongly with the ELISA responses (MHC-I *R_S_* = 0.86, MHC-II *R_S_* = 0.86), suggesting that the computational models can effectively guide vaccine design prior to experimental validation.

**Extended Data Fig. 7:**
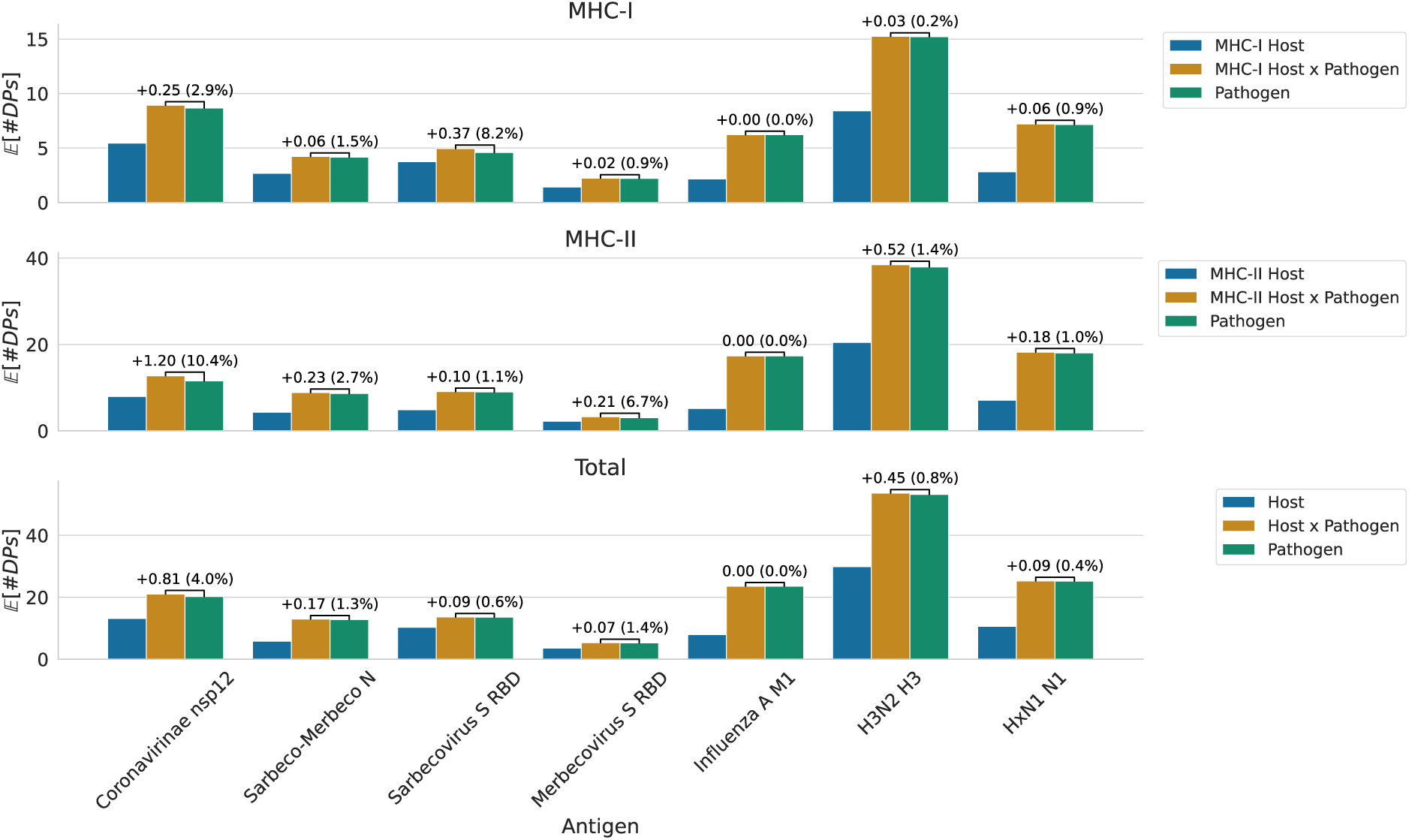
Attribution of host and pathogen coverage for the expected number of displayed peptides. For each of the seven *Coronavirinae* and Influenza A antigens (x-axis), three antigens were designed using Spectravax with different coverage score configurations: Host coverage only (blue), Pathogen coverage only (green), and the combined Host x Pathogen coverage (yellow). The expected number of displayed peptides (E[#DPs]) is shown on the y-axis. Each subplot represents the antigens designed and evaluated for MHC-I (**A**), MHC-II (**B**), and both MHC classes combined (**C**). The results indicate that the pathogen coverage score is the primary contributor to E[#DPs], as antigens designed using pathogen coverage alone exhibit higher values than those using host coverage alone. However, antigens designed with the combined Host x Pathogen coverage achieve the highest E[#DPs] across all antigens and MHC types. This demonstrates that incorporating host coverage further enhances the expected number of displayed peptides. The difference between the combined and pathogen-only designs, annotated on the figure, highlights the added value of including host coverage in antigen design, as implemented in Spectravax, compared to methods like Epigraph that consider only pathogen coverage.

**Extended Data Table 2:**
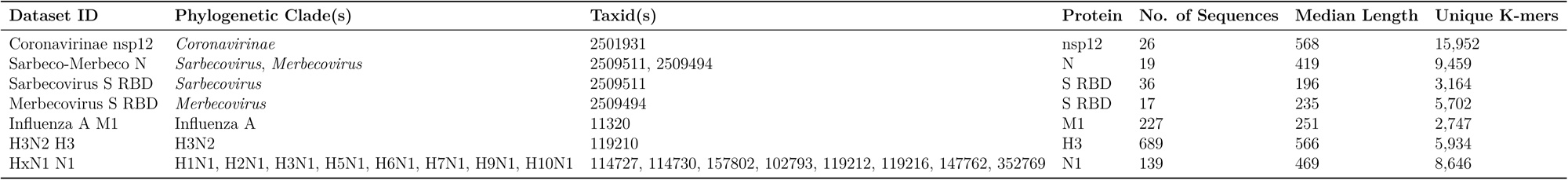
Datasets used to evaluate Spectravax. The following antigens were selected from within the coronavirus (*Coronavirinae*) subfamily: nucleoprotein (N), spike (S) and non-structural protein 12 (nsp12). RBD refers to the receptor-binding domain of the spike protein. Influenza A virus (within the *Alphainfluenzavirus influenzae* species) antigens include: hemagglutinin (H3), neuraminidase (N1) and matrix protein 1 (M1). To design a human-specific H3N2 vaccine, only sequences from the human host were downloaded. All numbers refer to the processed datasets.

**Extended Data Fig. 8:**
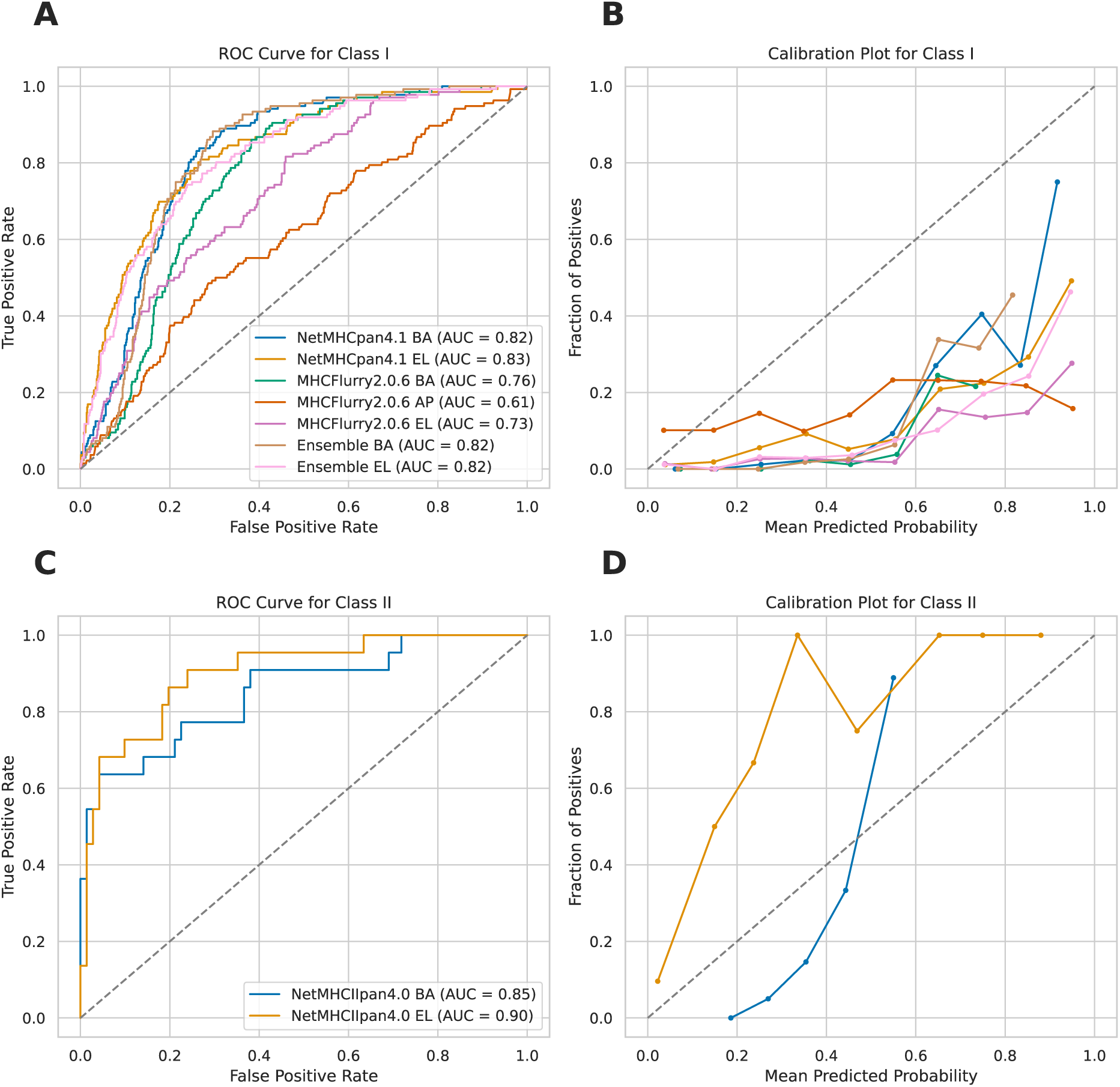
Performance of computational methods for predicting peptide-MHC binding. Evaluation of computational methods using SARS-CoV-2 peptide stability data across 11 MHC allotypes (5 HLA-A, 1 HLA-B, 4 HLA-C, 1 HLA-DRB1)^62^. The dataset includes 777 unique peptides, 174 of which were binders as determined by the NeoScreen assay. We assessed NetMHCpan v4.1, NetMHCIIpan v4.0^14^, and MHCflurry v2.0.6^63^, predicting Binding Affinity (BA), Antigen Processing (AP), and Eluted Ligand (EL) scores. (**A**, **C**) Receiver Operating Characteristic (ROC) curves for MHC-I and MHC-II, respectively, showing the trade-off between true positive rate (TPR) and false positive rate (FPR). The area under the ROC curve (AUROC) indicates model performance, with higher values representing better discrimination between binders and non-binders. (**B**, **D**) Calibration plots for MHC-I and MHC-II, respectively, assessing the reliability of predicted probabilities. Perfect calibration aligns with the diagonal dashed line. The NetMHCpan EL score outperformed other methods with AUROC of 0.83 (MHC-I) and 0.90 (MHC-II), and showed good calibration, supporting its use as the default criterion for predicted binding in our study.

**Extended Data Fig. 9:**
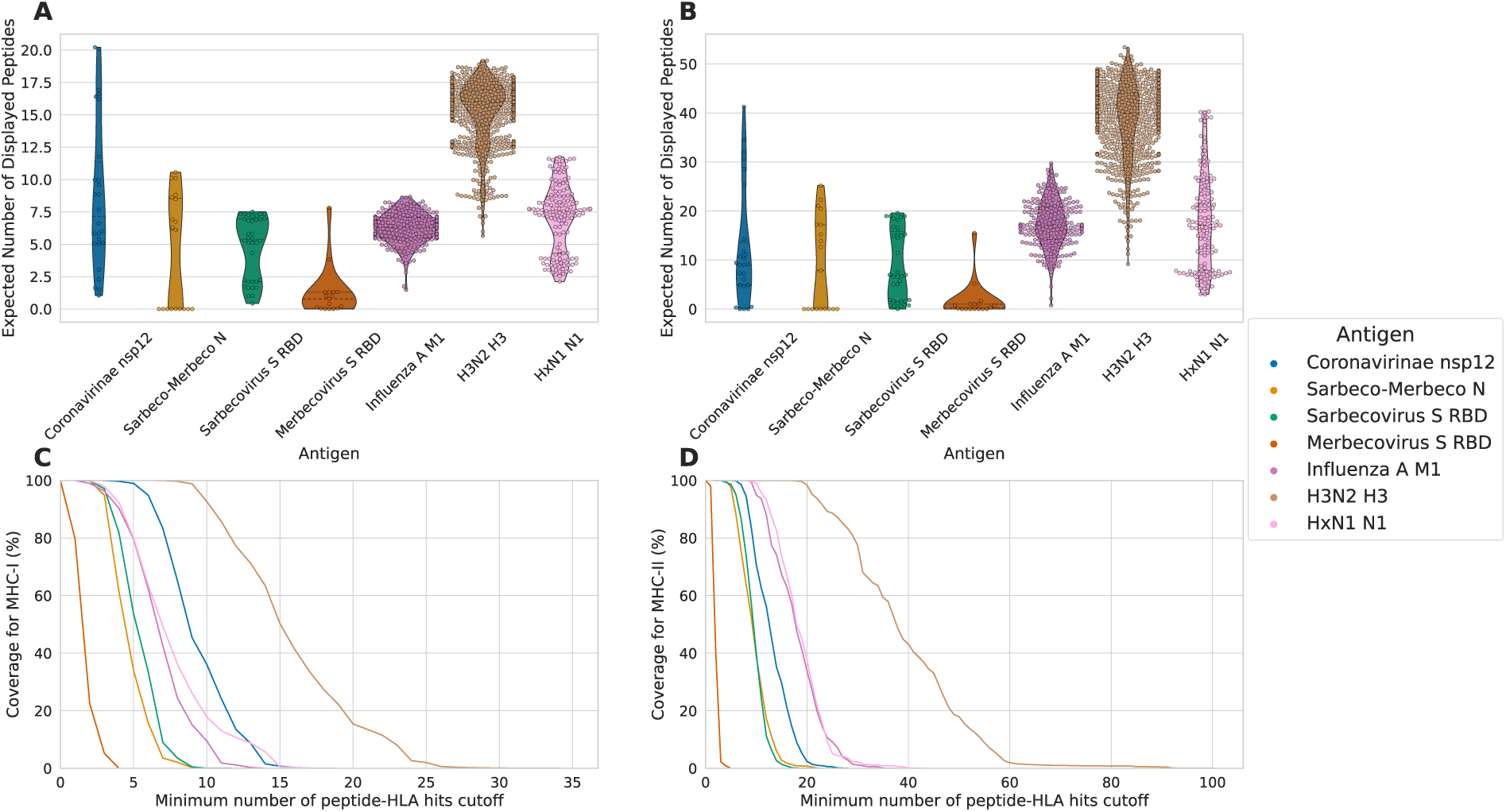
Comparison of the coverage of different Spectravax antigen designs. To demonstrate the generalisability of Spectravax, we applied the method to design six additional vaccine antigens from two diverse clades of viruses: Influenza A viruses and *Orthocoronavirinae* representing realistic targets for broad-spectrum vaccine design. All antigens were designed using default parameters (Extended Data Table 3). (**A, B**) Violin plots showing the E[#DPs] of the Spectravax antigen designs for (**A**) MHC-I and (**B**) MHC-II. The x-axis shows the different antigens and the y-axis shows the expected number of displayed peptides. Each point represents a different pathogen in the target sequences. (**C, D**) Line plots showing the coverage of the average host population for (**C**) MHC-I and (**D**) MHC-II. The x-axis shows the minimum number of peptide-HLA hits cutoff (*n*) and the y-axis shows the coverage, which is the fraction of the host population predicted to display at least *n* peptides.

**Extended Data Table 3:**
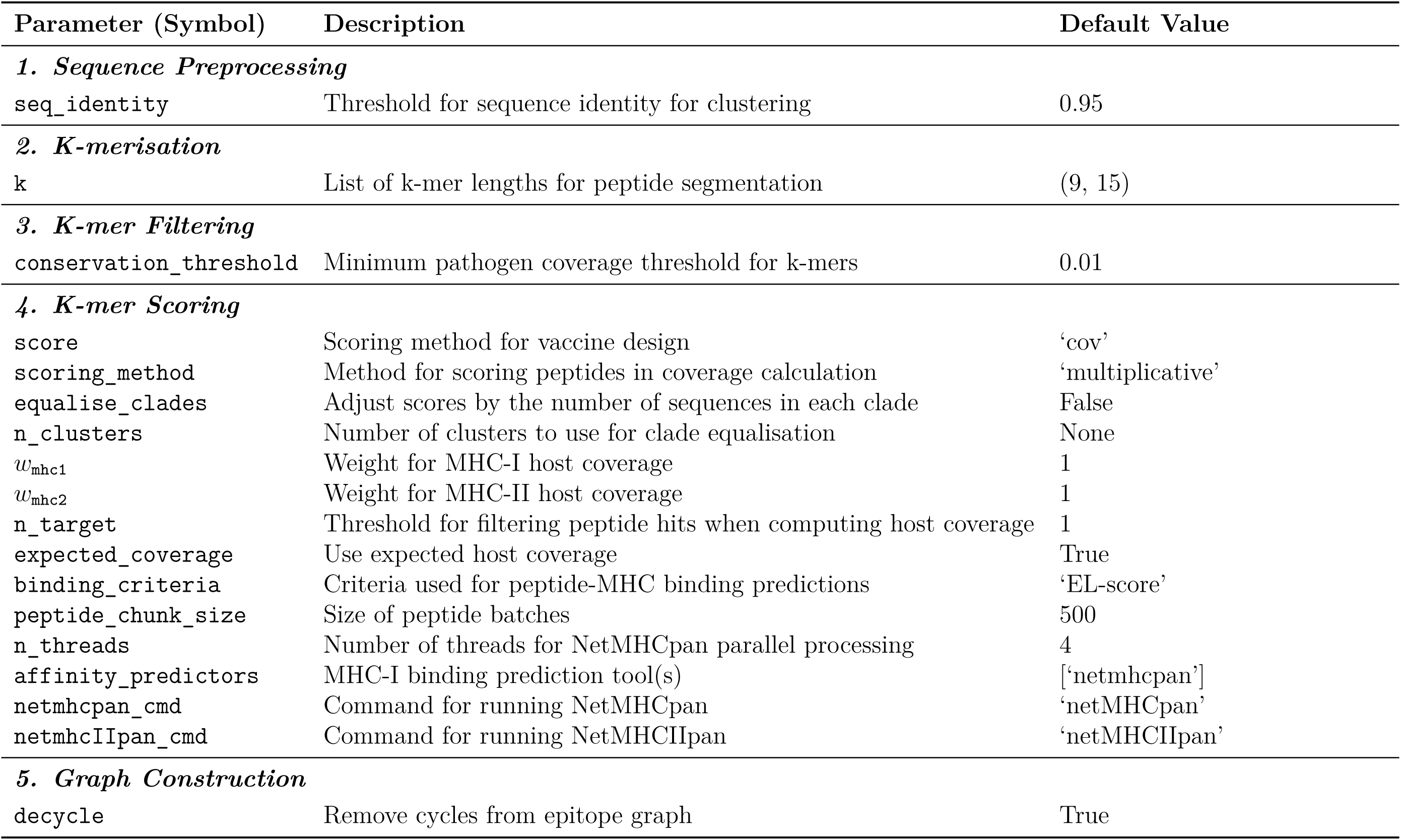
Default parameters used in the Spectravax pipeline.

**Exteded Data Fig. 10:**
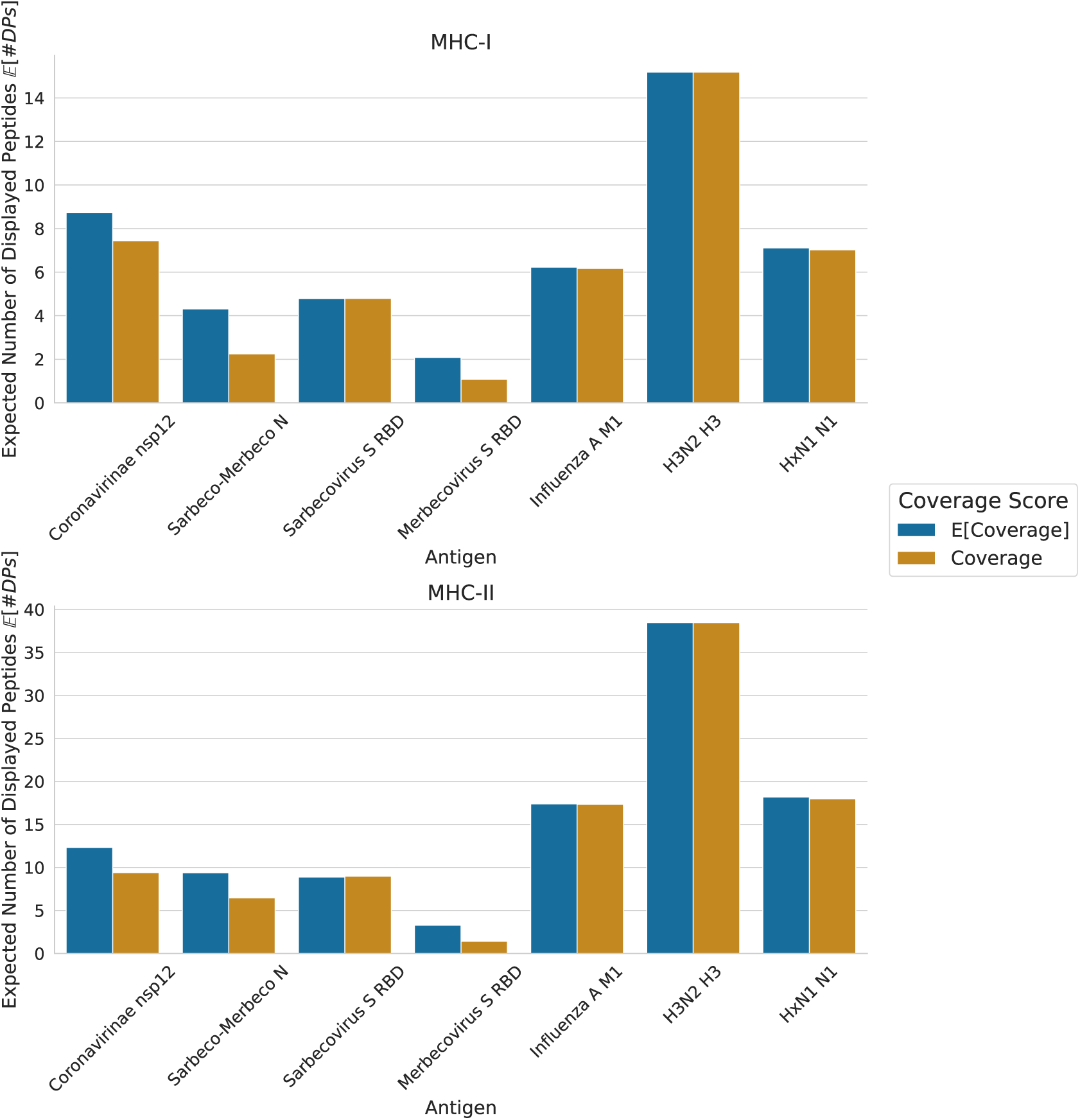
Comparison of antigens designed using the coverage and expected coverage criteria in terms of expected displayed peptides. For each of the seven *coronavirinae* and influenza A antigens (x-axis), two antigens were designed using Spectravax with the following coverage scores: E[Coverage] (blue) and Coverage (yellow). The expected number of displayed peptides (E[#DPs]) is shown on the y-axis. The subplots show evaluations for MHC-I and MHC-II.

